# Examining speech-brain tracking during early bidirectional, free-flowing caregiver-infant interactions

**DOI:** 10.1101/2024.05.16.594544

**Authors:** E.A.M. Phillips, L. Goupil, J. E. Ives, P. Labendzki, M. Whitehorn, I. Marriott Haresign, S.V. Wass

## Abstract

Neural entrainment to slow modulations in the amplitude envelope of infant-directed speech is thought to drive early language learning. Most previous research with infants examining speech-brain tracking has been conducted in controlled, experimental settings, which are far from the complex environments of everyday interactions. Whilst recent work has begun to investigate speech-brain tracking to naturalistic speech, this work has been conducted in semi-structured paradigms, where infants listen to live adult speakers, without engaging in free-flowing social interactions. Here, we test the applicability of mTRF modelling to measure speech-brain tracking in naturalistic and bidirectional free-play interactions of 9-12-month-olds with their caregivers. Using a backwards modelling approach, we test individual and generic training procedures, and examine the effects of data quantity and quality on model fitting. We show model fitting is most optimal using an individual approach, trained on continuous segments of interaction data. Corresponding to previous findings, individual models showed significant speech-brain tracking at delta modulation frequencies, but not in alpha and theta bands. These findings open new methods for studying the interpersonal micro-processes that support early language learning. In future work, it will be important to develop a mechanistic framework for understanding how our brains track naturalistic speech during infancy.

## 1. Introduction

Understanding how infants learn language from inconsistent and irregular environmental input is one of the key challenges in developmental neuroscience (Leong & Goswami, 2015; Yu et al., 2021; Yu & Smith, 2012). A large body of literature has shown that adults’ neural oscillations entrain to the amplitude modulations of continuous speech (Luo & Poeppel, 2007), and recent work is beginning to explore when and how the developing brain first begins to track these temporal modulation structures (e.g. Attaheri et al., 2022). However, so far, this work has been conducted in structured and controlled experimental paradigms which are far from the cluttered, dynamic and complex naturalistic settings in which infants begin to parse the continuous speech stream and learn to decode the phonology of their native language (Yu et al., 2021; Yu & Smith, 2012).

Neural tracking of the speech amplitude envelope at different temporal modulation frequencies supports our ability to parse the speech signal into linguistic units (Giraud & Poeppel, 2012; Gross et al., 2013; Leong et al., 2017; Poeppel, 2014). In adults, neural tracking has been shown at delta, theta and low gamma frequencies, corresponding to the stress, syllabic and phonetic patterning of the phonological information in speech (Ding et al., 2016; Gross et al., 2013; Leong & Goswami, 2015). Oscillatory tracking of a speech signal in the auditory cortex is thought to be hierarchically nested, with slower theta-rate oscillations dynamically modulating the activity of oscillations at higher frequencies (Giraud & Poeppel, 2012). In particular, the phase of theta-rate oscillations is thought to modulate the power and phase of gamma oscillations, supporting the integration of syllabic and phonetic information (e.g. Gross et al., 2013).

Whilst neural tracking of theta-rate modulations is thought particularly important for speech encoding and intelligibility in adults (Doelling et al., 2014), greater attention has been paid in infant research to the role of delta-rate modulations in early phonological development (Leong & Goswami, 2015). Recent work has, for example, shown that the peak modulation frequency of infant directed speech (IDS) – i.e. the modulation rate with the most energy - occurs at delta frequencies (∼2Hz; Leong et al., 2017). This is in contrast to ADS, where peak modulation rates occur within the theta band (∼5Hz; Greenberg et al., 2003; Leong et al., 2017). Given that delta-rate temporal modulations correspond to the timing and proportion of stressed syllables in continuous speech, Leong and colleagues have suggested that attunement to delta rate modulations could be particularly important in helping infants locate word boundaries, as well as grouping speech streams into phonological units during early language learning (Leong et al., 2017; Leong & Goswami, 2015; Power et al., 2013).

In adults, speech tracking has often been examined by recording the neural response of participants to short, repeated experimental stimuli. For example, one popular method involves presenting participants with a repeated speech stimulus, and calculating the inter-trial phase consistency across neural responses (Luo & Poeppel, 2007; Zion Golumbic et al., 2013). Alternatively, researchers compute the phase locking value (measuring the similarity in phase between two signals) between the amplitude envelope of a speech stimulus and participants’ neural responses at the same frequency, and average the phase-locking value across segments of short, repeated trials (Doelling et al., 2014; Peelle et al., 2013).

In more recent years, however, the development of novel methods of analysis to examine ongoing neural responses to a continuous speech signal has become a particular focus (Crosse et al., 2015, 2016, 2021; Jessen, 2019; Lalor et al., 2009; Obleser & Kayser, 2019). One popular approach is to model the impulse response function between the continuous speech signal and the ongoing EEG response (Brodbeck & Simon, 2020; Crosse et al., 2016, 2021; Ding & Simon, 2012). These models are referred to as multivariate temporal response functions (mTRFs), that estimate the linear mapping from an environmental stimulus to the EEG response via a method of regularised regression (Crosse et al., 2016). Through an iterative process of training and testing, a regression model is computed at each electrode, over a range of time-lags (Crosse et al., 2016), to describe the relationship between the speech amplitude envelope (e.g. 1-8Hz; Kalashnikova et al., 2018) and the EEG signal. The predictive accuracy of the model is then computed by calculating the Pearson’s correlation between the actual neural response and that predicted by the model (Crosse et al., 2016).

As well as forwards (encoding) models, that map from the stimulus to the EEG response, backwards (decoding) mTRF models can also be used to model the linear mapping from the stimulus (speech signal) back to the neural response. This method is widely used to investigate speech-brain tracking in the adult literature (Bednar & Lalor, 2020; Crosse et al., 2015; Ding & Simon, 2012). Unlike forward (encoding) models, backwards models do not require the pre-selection of electrodes to enter into the model. Instead, during model training, each channel is assigned weights at each time lag, based on how much information that channel provides for the reconstruction. In this way, channels that track the speech signal to a very little extent and those that track the signal to a great extent will both have high weights because they provide most information to the reconstruction (Crosse et al., 2016).

Two different approaches to model training can be taken for forwards and backwards models: an individual training approach, or a generic approach. Individual model training is the most commonly used method, whereby an mTRF model is trained on the trials of an individual participants’ data, and then tested on held aside data for that participant. In this way, individual training methods identify consistencies in neural tracking within participants (Jessen, 2019; Jessen et al., 2021). The second approach is a generic training approach: here, an mTRF model for each participant is trained by pooling across the models constructed for all other participants. This method is considered more optimal with small or noisier data sets because using data across all participants increases the amount of data the model is trained on comparative to the individual training approach (Crosse et al., 2021; Jessen et al., 2021). In training models across participants, generic models also inform our understanding of consistencies in neural tracking of the speech signal between participants (Jessen et al., 2021), with higher predictive accuracies suggesting greater model consistencies between participants. This is particularly interesting for examining speech-brain tracking in naturalistic interactions where each infant listens to a different speech signal (i.e. their caregivers’).

Three studies have recently employed mTRF modelling methods to examine audio-brain tracking in early infancy. Jessen and colleagues used a forwards modelling generic training approach to compute an mTRF model of the relationship between 11-12-month-old infants’ broadband EEG signal (1-40Hz) and the amplitude envelope of a cartoon video that each infant engaged with for just over two minutes. They identified significant speech-brain tracking of the amplitude envelope over fronto-central electrodes, which was also present, but with greater inter-participant variability, using an individual modelling procedure (Jessen, 2019). Presenting infants with pre-recorded, naturalistic infant-directed speech, recorded from a caregiver interacting with their infant, Kalashnikova et al. (2018) showed significant speech-brain tracking by infants to the amplitude envelope (filtered between 0.5-8Hz), using an individual training, forwards mTRF modelling procedure. Where infants were instead presented with the same speaker, speaking in adult-directed speech, no speech-brain tracking was observed.

More recently, Attaheri et al. (2022) constructed a backwards mTRF to model the relationship between 4, 7 and 11-month-old infants’ continuous neural activity and sung nursery rhymes. The pre-recorded screen-based stimuli used in their study consisted of a British female speaker melodically singing or chanting nursery rhyme phrases, with a mean length of 4s. To ensure consistent beat rates across all nursery rhymes, the speaker listened to a 120 bpm metronome. Individual training models were computed between the amplitude envelope of the nursery rhyme phrases (filtered at 0.5-15Hz), and infants’ EEG responses, filtered into three frequency bands: delta (0.5-4Hz), theta (4-8Hz) and alpha (8-12Hz).

Corresponding to the prediction that infants track the amplitude envelope of speech at low modulation frequencies (Leong et al., 2017), across all 3 ages, significant speech-brain tracking was identified at delta and theta frequencies but not alpha frequencies.

Almost all previous work examining neural tracking to the temporal modulation structure of speech in infant and adult populations has, however, been conducted using structured experimental paradigms, where participants are presented with a continuous speech sound, and their neural response to that speech sound is recorded (Attaheri et al., 2022; Bednar & Lalor, 2020; Crosse et al., 2015; Jessen, 2019; Kalashnikova et al., 2018). This is very different to the speech that infants hear in naturalistic, everyday settings where vocalisations are variable in length, often repetitive and short, and occur in the context of background noise (Gratier et al., 2015; Henning et al., 2005). In very early face-to-face interactions, with infants aged 8-13 weeks, for example, caregiver vocalisations have been found to last an average of 1s, and range between 0.5-9s (Gratier et al., 2015; Jaffe et al., 2001). In the home environment, face-to-face, joint engagement interactions tend to occur in bursts throughout the day (Suarez-Rivera et al., 2022), and recent work, recording from caregivers and infants in their homes over day-long time periods, has shown that the organisation and content of these face-to-face interactions, are particularly important to early word learning (Custode & Tamis-LeMonda, 2020; Schatz et al., 2022). Examining speech-brain tracking during naturalistic free-flowing interactions is also important to informing our understanding of how infant processing of the temporal modulation structure of speech is related to the ongoing dynamics of shared interactions (joint attention behaviours, for example), as well as its relationship to the semantic timing and complexity of the caregivers’ speech inputs (Nencheva & Lew-Williams, 2022).

Examining speech-brain tracking during naturalistic interactions is, however, particularly problematic given the noise inherent to both the infants’ EEG signal and the caregivers’ speech (Georgieva et al., 2020; Noreika et al., 2020). Naturalistic EEG recordings are particularly affected by movement artifact, and although ICA decomposition removes some of these components, it does not remove all (Marriott Haresign et al., 2021). During shared object play, the caregiver’s speech signal is also affected by background noise in the interaction, including toy clacks, as well as infant movement and co-vocalisations (Gratier et al., 2015).

Two recent studies have made some advances in examining speech-brain tracking in more naturalistic, semi-structured paradigms. Menn and colleagues examined the correlation between infants’ EEG activity and their caregiver’s speech signal, during a semi-structured paradigm, where the caregiver described objects to their infant, but the infant was unable to touch or engage in play with the objects (Menn et al., 2022). Coherence between infant EEG activity and the caregivers’ speech signal was above chance for both syllabic and stress-rate amplitude modulations. Modelling the forwards mTRF response between natural caregiver singing and infant EEG activity at 1-10 Hz, Nguyen et al. (2020) also identified speech brain tracking above levels predicted by chance, among 7-month-old infants. In their paradigm, infants watched cartoon videos on an iPad positioned on their caregiver’s lap, whilst listening to the caregivers’ sung speech.

Both studies have, therefore, examined speech-brain tracking during naturalistic, but uni-directional interactions (i.e. where the caregiver is speaking or singing to the infant, and the infant is passively listening to the caregivers’ speech inputs). Though such paradigms reduce the amount of movement artifact in the infant’s EEG signal, as well as background noises such as toy clacks and infant vocalisations that introduce noise to the caregivers’ speech signal, the structured nature of the paradigms limits our understanding of speech-brain tracking in everyday, bidirectional interactions, where caregivers and infants take turns in leading and responding to vocal and joint-attentional cues in their partner. Do infants track the slow amplitude modulation patterns of their caregiver’s speech during dynamic and free-flowing interactions? And, at what time-scales do we observe neural tracking, given the complexities in the timing and length of the caregivers’ vocalisations in naturalistic interactions? Examining these questions is crucial to furthering our understanding of the neural and behavioural processes that support early language development.

Here, we aim to examine speech-brain tracking during naturalistic, interactive caregiver-infant interactions using a backwards mTRF modelling approach. To test whether patterns of speech-brain tracking that are observed in controlled experimental paradigms transfer to interactive contexts, we match the analysis design of (Attaheri et al., 2022). In particular, we compute mTRF models that map the wideband amplitude envelope (0.5-15Hz) to the infant EEG signal at delta (1-4Hz), theta (4-8Hz) and alpha (8-12Hz) frequencies to test whether speech-brain tracking differs significantly from chance at slow modulation frequencies only.

Given that this is the first time that continuous methods have been used to examine speech-brain tracking during naturalistic play, we take two different approaches to segmenting the caregivers’ speech signal as inputs to the mTRF model. First, we train the mTRF models on continuous data segments, dividing each interaction up into 10 equal-length parts, corresponding to the approach most often taken in experimental designs with adults, and infants, where a continuous stimulus is presented (Jessen, 2019). This approach also optimises the amount of data on which the model was trained. Data quality and quantity have been shown to affect the performance of mTRF models, such that noisier data, and models trained on small data sets, often lead to overfitting, resulting in poor transferability to new data (Crosse et al., 2021). However, due to the fact that, in adult studies, data tends to be more plentiful and of higher quality, extensive testing of how data quality and quantity influence model performance has not been conducted (Mesik & Wojtczak, 2022).

Second, we computed mTRF models with each caregiver vocalisation serving as an individual fold in the model (see Sohoglu and Davis (2020) for a similar approach with adults). Though this approach decreased the amount of data that each individual model is trained on, it also reduced the amount of noise in the data (i.e. in this way only periods that the caregiver is speaking are included in the model, and this also means that analyses can be run on vocalisations where no background noise or co-vocalisations occurred within a vocalisation). Training the model on individual vocalisations also gives us the opportunity to examine the effect of vocalisation length on speech-brain tracking. As mentioned above, during naturalistic joint attentional interactions, caregiver vocalisations tend to be short, in contrast to the continuous speech streams presented to infants in controlled and semi-naturalistic designs. Here, we test whether speech-brain tracking is greater, or observed only, for longer vocalisations (lasting 2000ms or longer), in comparison to models trained on all vocalisations lasting over 500ms. We therefore computed separate mTRF models for all vocalisations lasting 500ms or longer, vocalisations lasting 2000ms or longer, as well as vocalisations lasting 500ms or longer, or 2000ms or longer that included no background noise (e.g. toy clacks, and infant vocalisations).

Given that inputting individual vocalisations into the model reduces the amount of data on which the models are trained (particularly where only longer (> 200ms) or clean vocalisations are included), we tested for overfitting of the mTRF models to the training data. To do so, we compare the predictive accuracy scores obtained during model training and testing to examine whether the accuracy scores for training sets, computed during the cross-validation procedure, are higher than for testing sets (see Methods (section 2) for more details). If so, this would indicate poor transferability to new data, and therefore insufficient model fitting (Crosse et al., 2021). These comparisons are contrasted with the continuous methods of data chunking that optimise the amount of data entered into the model.

In contrast to Attaheri et al. (2022), where an individual training method alone was used, we also present models computed according to generic training procedures. This is due to the noisiness of our data, and the reduced amount of data entered into the vocal chunking models. As mentioned above, in pooling data across participants during model training, generic models maximise the amount of data on which the mTRF model is computed, in comparison to individual models which use the data of just one participant at a time. As well as being methodologically beneficial, generic models also provide an interesting comparison to the individual models in testing for consistencies in speech tracking by infants to their own caregiver’s speech signal across participants (Jessen, 2019). This has, to our knowledge, not previously been examined.

Based on the findings of the recent experimental paradigms conducted with infants, we hypothesised that speech-brain tracking would be observed at slow amplitude modulation frequencies (delta and theta) for both individual and generic models (Attaheri et al., 2022; Jessen et al., 2021). Considering the comparison of models trained on continuous auditory streams vs. individual vocalisation chunks, it was hypothesised that speech-brain tracking would be observed across all models: those trained on continuous data streams, models trained on vocalisations lasting 2000ms or more and vocalisations lasting 500ms or more (Attaheri et al., 2022). Given the susceptibility of mTRF models of over-fitting to the training data where smaller training data sets are used, we hypothesised that models trained on continuous data segments would be associated with increased speech-brain tracking values, compared to the vocalisation chunk models, given that much more data would be used to train the continuous models. Finally, based on previous comparisons of audio-brain tracking using individual vs. generic model training methods with data collected from infants watching cartoon videos (Jessen, 2019), it was hypothesised that greater speech-brain tracking would be observed where mTRF models were computed using individual training methods.

## 2. Method

### 2.1 Participants, equipment and pre-processing

#### 2.1.1. Participants

Ninety-four caregiver-infant dyads took part in this study. The final overall sample with usable, coded, gaze data was 46 (32 infants were excluded due to recording error or equipment failure, 4 infants were excluded for fussiness and 12 infants were excluded due to poor quality EEG data). For the final, overall sample, the mean age of participants was 10.93 months (SD=1.29; 26 females, 20 males). All caregivers were female. Participants were recruited through baby groups and Children’s Centers in the Boroughs of Newham and Tower Hamlets, as well as through online platforms such as Facebook, Twitter and Instagram. Written informed consent was obtained from all participants before taking part in the study, and consent to publish was obtained for all identifiable images used. All experimental procedures were reviewed and approved by the University of East London Ethics Committee.

#### 2.1.2. Experimental set-up

Parents and infants were seated facing each other on opposite sides of a 65cm wide table. Infants were seated in a high-chair, within easy reach of the toys (see Figure 1b). The shared toy play comprised two sections (play section 1 and play section 2), with a different set of toys in each section, each lasting ∼5 minutes each. Two different sets of three small, age-appropriate toys were used in each section; this number was chosen to encourage caregiver and infant attention to move between the objects, whilst leaving the table uncluttered enough for caregiver and infant gaze behaviour to be accurately recorded (cf. Yu & Smith, 2013).

**Figure 1.**
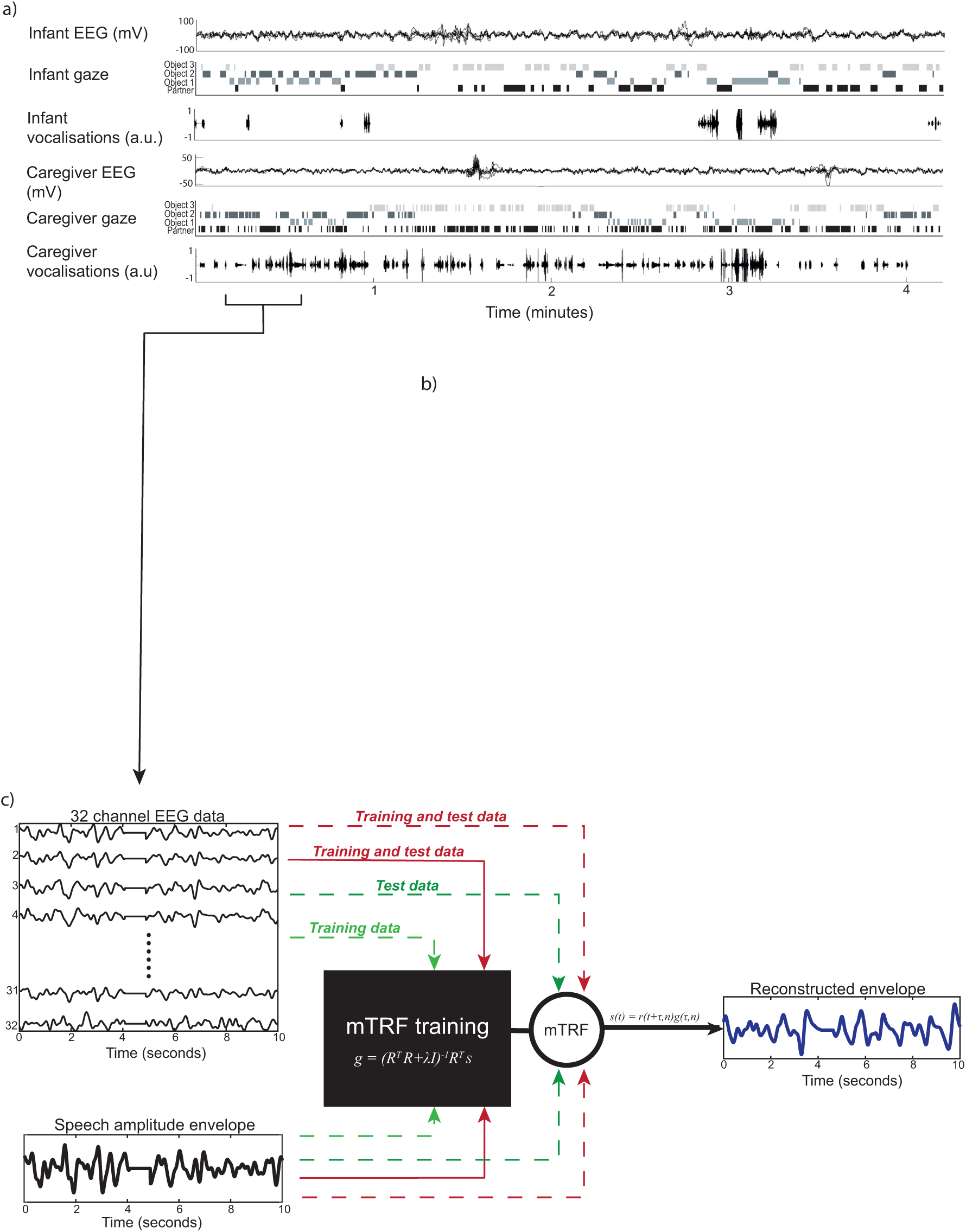
Experimental set-up and mTRF computation. a) Raw data sample, showing (from top) infant EEG over fronto-central electrodes, after pre-processing, infant gaze behaviour, infant vocalisations, adult EEG over fronto-central electrodes, adult gaze behaviour, adult vocalisations. b) Example camera angles for caregiver (top and middle) and infant (bottom), showing right and left cameras synchronized to the caregiver camera (middle). c) Depiction of mTRF model computation. Left panel shows example EEG data (top), filtered at delta frequencies only, and the speech amplitude envelope (bottom), after pre-processing and z-scoring, for one participant. Middle panel shows how data were partitioned before being submitted to model training and testing for the generic (red) and individual (green) models. Dashed lines indicate data inputs for individual participants, whilst solid lines indicate data inputs across all participants. Right panel shows the reconstructed amplitude envelope, computed from testing data, for the individual model only.

At the beginning of each play section, a researcher placed the toys on the table, in the same order for each participant, and asked the caregiver to play with their infant just as they would at home. Both researchers stayed behind a screen out of view of caregiver and infant, except for the short break between play sessions.

#### 2.1.3. Equipment

EEG signals were recorded using a 32-channel BioSemi gel-based ActiveTwo system with a sampling rate of 512Hz with no online filtering using Actiview Software. The interaction was filmed using three Canon LEGRIA HF R806 camcorders recording at 50 fps. Parent and infant vocalisations were also recorded throughout the play session, using a ZOOM H4n Pro Handy Recorder and Sennheiner EW 112P G4-R receiver.

Two cameras faced the infant. One was placed on the left of the caregiver, and one on the right (see Figure 1b). Cameras were placed so that the infant’s gaze and the three objects placed on the table were clearly visible, as well as a side-view of the caregiver’s torso and head. One camera faced the caregiver, positioned just behind the left or right side of the infant’s high-chair (counter-balanced across participants). One microphone was attached to the caregiver’s clothing and the other to the infant’s high-chair.

Caregiver and infant cameras were synchronised to the EEG via radio frequency (RF) receiver LED boxes attached to each camera. The RF boxes simultaneously received trigger signals from a single source (computer running MATLAB) at the beginning of each play section, and concurrently emitted light impulses, visible in each camera. Microphone data was synchronised with the infants’ video stream via a xylophone tone recorded in the infant camera and both microphones, which was hand identified in the recordings by trained coders. All systems were extensively tested and found to be free of latency and drift between EEG, camera and microphone to an accuracy of +/- 20 ms.

All caregivers and infants also took part in two further interactions, including a caregiver puppet show interaction and a section where infants played with toys on their own. These interactions are, however, not included in the current analyses.

#### 2.1.4. Vocalisation coding

The onset and offset times of caregiver and infant vocalisations were first provisionally identified using an automatic detector (previously used in Phillips et al., accepted). The algorithm detected voiced segments and compared the volume and fundamental frequency in each recorded channel to infer the probable speaker (mother vs. infant). Identification of the onset and offset times of the detector then underwent a secondary analysis by trained coders, who also identified misidentification of utterances by the automatic decoder, and classified the speaker for each vocalisation. We also identified that the detector did not accurately identify onset and offset times of caregiver and infant speech during co-vocalisations.

Because vocalisations by the infant during the caregivers’ speech would introduce noise to the caregivers’ speech signal, as well as the possibility that infants’ were entraining to their own speech (Pérez et al., 2021), vocalisations that contained overlapping speech segments were therefore excluded from the vocal chunking analysis. Initially, the detector was programmed so that a gap of 250ms was placed between vocalisations (i.e., if there was a pause of >250ms, then the utterance was treated as two separate vocalisations). After hand-coding, the vocalisations were re-processed so that a 1000ms gap was placed between each vocalisation. This step was taken to increase the lengths of the vocalisations included in the vocal chunking analysis (Leong et al., 2017).

In addition to the hand-coding of the vocalisations based on the microphone data, the gaze behaviours of caregivers and infants, recorded in the 3 cameras, were also hand-coded throughout each interaction to a time resolution of 50Hz. Periods were marked as uncodable in the infant’s gaze time-series where the experimenter was present in the line of sight of either of the participants and/ or where it was unclear where the infant was looking. In order to exclude periods where the experimenter was present, and possibly speaking, all periods marked as uncodable were excluded from analysis.

#### 2.1.5. Infant EEG artifact rejection and pre-processing

A fully automatic artifact rejection procedure including ICA was adopted, following procedures from commonly used toolboxes for EEG pre-processing in adults (Bigdely-Shamlo et al., 2015; Mullen, 2012) and infants (Debnath et al., 2020; Gabard-Durnam et al., 2018), and optimised and tested for use with our naturalistic infant EEG data (Georgieva et al., 2020; Marriott Haresign et al., 2021). This was composed of the following steps: first, EEG data were high-pass filtered at 1Hz (FIR filter with a Hamming window applied: order 3381 and 0.25/ 25% transition slope, passband edge of 1Hz and a cut-off frequency at -6dB of 0.75Hz). Although there is debate over the appropriateness of high pass filters when measuring ERPs (see Widmann & Schröger, 2012), previous work suggests that this approach obtains the best possible ICA decomposition with our data (Dimigen, 2020; Marriott Haresign et al., 2021). Second, line noise was eliminated using the EEGLAB (Bigdely-Shamlo et al., 2015) function *clean_line.m* (Mullen, 2012).

Third, the data were referenced to a robust average reference (Bigdely-Shamlo et al., 2015). The robust reference was obtained by rejecting channels using the EEGLAB *clean_channels.m* function with the default settings and averaging the remaining channels. Fourth, noisy channels were rejected, using the EEGLAB function *clean_channels.m.* The function input parameters ‘correlation threshold’ and ‘noise threshold’ (inputs one and two) were set at 0.7 and 3 respectively; all other input parameters were set at their default values. Fifth, the channels identified in the previous stage were interpolated back, using the EEGLAB function *eeg_interp.m*. Interpolation is commonly carried out either before or after ICA cleaning but, in general, has been shown to make little difference to the overall decomposition (Delorme & Makeig, 2004). Infants with over 21% (7) electrodes interpolated were excluded from analysis. After exclusion, the mean number of electrodes interpolated for infants was 0.244 (*SD*=1.11) for play section 1, and 3.12 (*SD*=2.16) for play section 2.

Sixth, the data were low-pass filtered at 20Hz, again using an FIR filter with a Hamming window applied identically to the high-pass filter. Seventh, continuous data were automatically rejected in a sliding 1s epoch based on the percentage of channels (set here at 70% of channels) that exceed 5 standard deviations of the mean channel EEG power. For example, if more than 70% of channels in each 1-sec epoch exceed 5 times the standard deviation of the mean power for all channels then this epoch is marked for rejection. This step was applied very coarsely to remove only the very worst sections of data (where almost all channels were affected), which can arise during times when infants fuss or pull the caps. This step was applied at this point in the pipeline so that these sections of data were not inputted into the ICA. The mean percentage of data removed in play section 1 was 13.433 (SD=16.617), and 5.11(SD=7.03) for play section 2.

Data collected from the entire course of the recording session were then concatenated and ICAs were computed on the continuous data using the EEGLAB function runica.m. This included play section 1 and play section 2, as well as two further five-minute toy-play interactions, conducted as part of the recording session, that are not included in the current analysis. These additional interactions were included to increase the amount of data submitted to the ICA. The mean percentage of ICA components rejected was 51.90% (SD=16.92). After ICA rejection, data from each play section were re-split. For representative examples of artifactual ICA components identified in our naturalistic EEG data, see (Marriott Haresign et al., 2021).

### 2.2. mTRF analysis

#### 2.2.1. Step 1: Pre-processing

##### Removal of clipped segments from the speech signal

Due to the caregiver being within variable distance of their own microphones, some saturation was identified in a sample of the microphone recordings. A stringent clipping identification algorithm was used to remove parts of the microphone data where clipping occurred (Hansen et al., 2021). Parts of the signal for which clipping was identified for longer than a period of 1ms were set as missing data points. The percentage of the speech signal set as missing data points at this stage was very low for all interactions: M=0.036% (SD=0.11) for play section 1 and M=0.10% (SD=0.55) for play section 2.

##### Amplitude envelope of the speech signal

The amplitude envelope of the caregivers’ speech signal, recorded from their microphone during the interaction, was extracted and filtered below 15Hz. First, the signal was band-pass filtered (using a 3^rd^ order Butterworth filter (forwards and backward)) into 9 frequency bands, with equal spacing along the cochlear, according to Greenwood’s (1990) equation. The absolute value of the analytic signal generated by the Hilbert transform was then computed and averaged across frequency bands (Gross et al., 2013). Next, following Attaheri et al. (2022), the resulting wholeband amplitude envelope was filtered between 0.5 and 15Hz as follows: low pass filter (5^th^ order Butterworth filter forwards and backwards); high-pass filter (3^rd^ order Butterworth filter forwards and backwards; based on Di Liberto et al., 2015). Finally, at this point, the speech signal was downsampled to the sampling rate of the EEG signal (512Hz).

##### EEG data

After pre-processing, the EEG signal, for each participant, was filtered into delta (1-4Hz), theta (4-8Hz) and alpha (8-12Hz) frequencies (Attaheri et al., 2022). First, missing data points were excluded from the time series. Next, the EEG signal was filtered into the 3 frequency bands of interest using the *pop_eegfiltnew* function in the EEGLab toolbox(Delorme & Makeig, 2004), using zero-phase bandpass Hamming-windowed FIR filters.

#### 2.2.2. Step 2: Separating data into individual folds for training and testing

##### Continuous chunking analysis

For the continuous analysis, the filtered EEG and speech streams for each interaction (play section 1 and play section 2), for each participant, were split into equal-length folds, and allotted into training and test sets (see Figure 2 for a schematic depiction). First, the filtered EEG signal and the amplitude envelope of the caregivers’ speech were both synchronised to the video-camera time-series, using the methods described above. Where one signal had been recorded longer than the other, that signal was shortened so that both data streams matched in length. The start and end video frames of each interaction were identified, and the beginning and end of the signals were cut off where necessary. The start of each interaction was identified from the xylophone ding in the infants’ camera, whilst the end of the interaction was taken as the infants’ penultimate look of the interaction.

**Figure 2.**
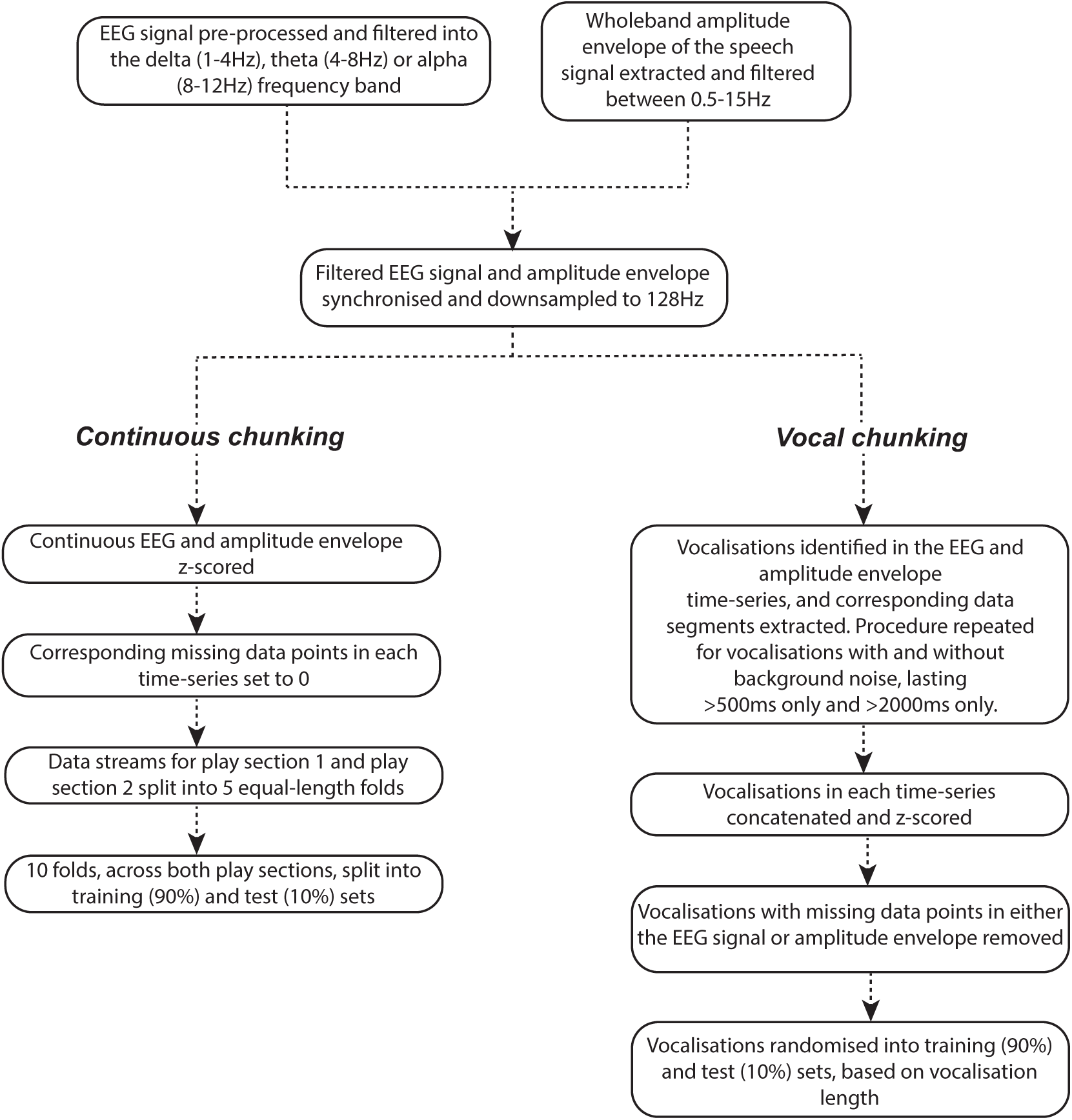
Schematic representation of the pre-processing steps to split data into individual folds for model training and testing. Procedures for the continuous chunking analysis (left) and vocal chunking analysis (right) are depicted.

Next, corresponding parts of the EEG (due to missing data) and speech signals (due to clipping) that were set as missing data points in one stream were identified in the other alternate stream and set to missing data points. To increase computational efficiency, both the EEG and speech streams were downsampled to 128Hz. These continuous data streams were then saved for permutation testing (see section 2.2.4 below). After each stream had been saved, both the speech and EEG streams were z-scored (EEG data were z-scored across channels), a step that is recommended when computing generic models for more consistent tuning of model parameters across data sets (Crosse et al., 2021). Finally, corresponding missing data points in both data streams were set to 0 (similar to Jessen et al., 2021).

The speech and EEG signals for play section 1 and play section 2 were then split into 5 equal-length folds, and randomised before one set was placed into the testing set. So that 90% of the folds were allotted to the training sets and 10% to the test set for each participant, one of the testing sets (from play section 1 or play section 2) was placed randomly into the training sets. Given that play section 1 and play section 2 were of different lengths, the combined training folds also varied in length (considered acceptable when training and optimizing mTRF models (Crosse et al., 2016). Where a participant had data for only play section 1 or play section 2 (which occurred where, for example, the infants’ EEG data was over the acceptable number of interpolated channels for one interaction), one fold was kept as the test set and the rest as the training sets. This step was taken to ensure that folds for each participant were of similar lengths.

##### Vocal chunking analysis

For the vocal chunking analysis, each caregiver vocalisation was entered into the mTRF model as an individual fold. First, each caregiver vocalisation was cut out of the EEG and speech time-series and treated as a separate data fold (see Figure 2 for a schematic depiction). First, the start and end of each caregiver vocalisation was identified in the synchronised time-series described above (up to the point that the data was downsampled to 128Hz) and cut out of the corresponding time-series. Where any missing data points occurred in either the EEG or speech segment for one vocalisation (i.e. those vocalisations with 1 missing data sample or more), this vocalisation was removed from analysis. Next, the periods of vocalisation identified in both time-series were concatenated and z-scored, and then converted back into discrete vocalisations for both the EEG and speech signals. Finally, vocalisations for each play section were randomised and allotted to training and test sets, so that 90% went into the training set and 10% into the test set. Our procedure for doing this grouped together vocalisations of similar lengths, so that vocalisations in the training and test sets were of similar sizes. For participants with play section 1 and play section 2, training and test sets from each play section were concatenated, randomised and adjusted to ensure that 10% of the vocalisations were in the test set and 90% in the training set. The procedure was repeated for vocalisations with and without background noise lasting over 500ms and over 2000ms, resulting in four separate sets of vocalisation chunks. Where caregivers had fewer than 5 vocalisations for a respective set, that dyad was excluded from the analysis.

#### 2.2.3. Step 3 - mTRF computation

mTRF analysis was carried out using the mTRF toolbox (Cross et al., 2016). As described in the introduction, in backwards mTRF modelling, all EEG channels are entered into the model at once, and assigned weights through the iterative regression modelling process, based on their degree of involvement in tracking the speech signal. The backwards mTRF model can be expressed by the following equation, where the TRF (*g*) represents the linear mapping from the neural response *r*(*τ*, *n*) back to the speech stimulus *ś*(*t*):

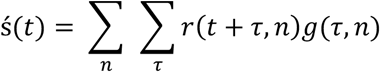

where, ś(*t*) is the estimated stimulus envelope, *g*(*τ*, *n*) is the TRF, and *r*(*τ*, *n*) is the EEG signal, over each time lag (*τ*), at each channel (*n*) included in the model. The TRF (*g*) is estimated using a method of regularized regression:

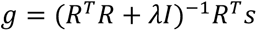

where *R* is the EEG data at each time lag, and *ś* is the zero-lagged stimulus. *R*^$^*R* is the autocovariance matrix of the neural response which is divided out from the model. In dividing out the autocovariance of the EEG response from the model, inter-channel redundancies are no longer included in the model, and, as a result, each TRF weight at each channel represents the amount of information that weight provides for the re-construction (Crosse et al., 2021). Both individual and generic models were computed over time-lags from -50 to 250ms (similar to (Attaheri et al., 2022).

##### Model training and testing

Models were trained using either an individual (subject-dependent) approach, or a generic modelling (subject-independent) approach. The analysis steps for each training approach are detailed below and depicted in Figures 1 and 3.

**Figure 3.**
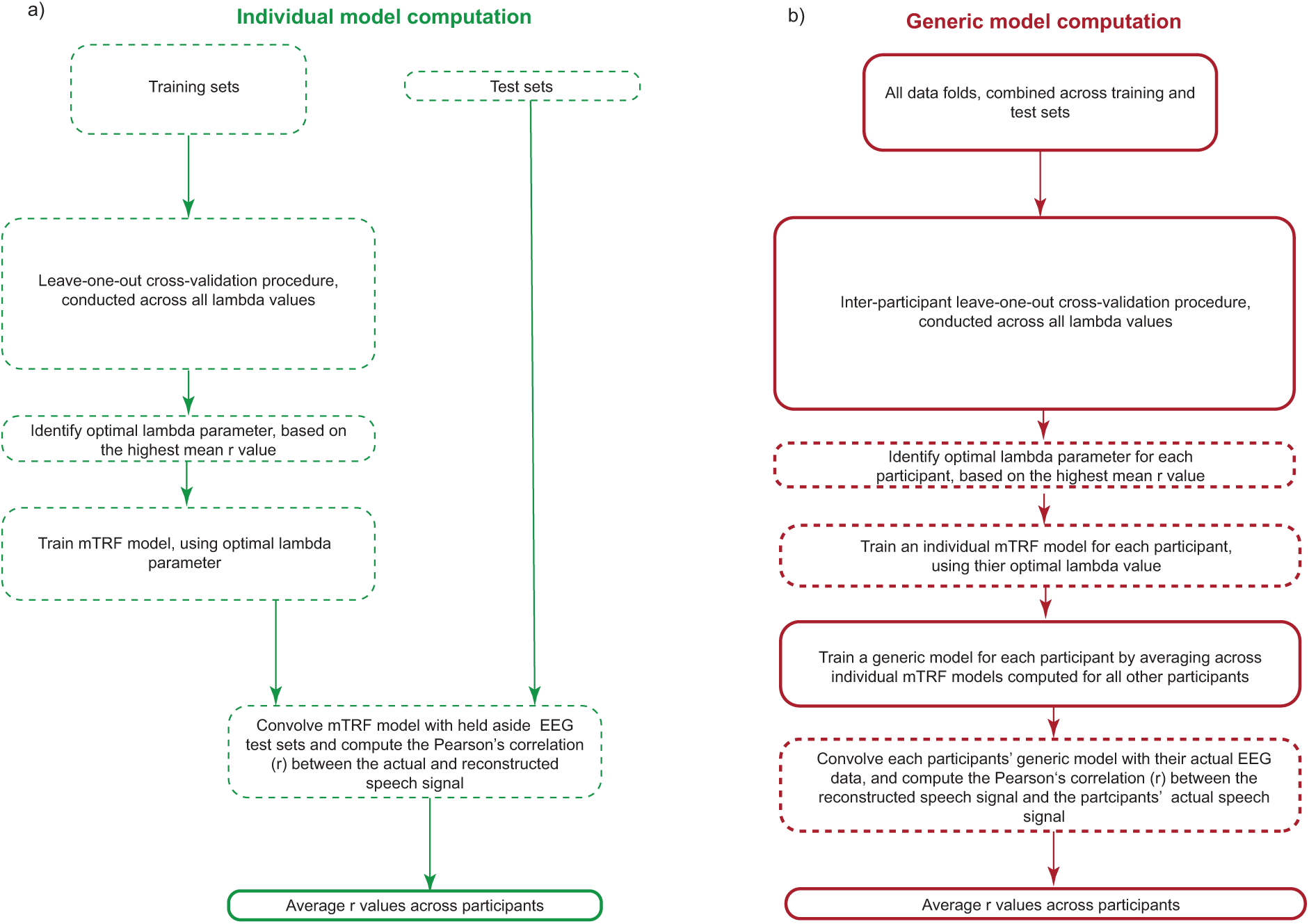
Schematic representation of mTRF model computation. a) Individual model computation. b) Generic model computation. Dashed boxes indicate procedures conducted on the data of each individual participant; hard-line boxes indicate procedures conducted across the data of all participants. The proportional size of the boxes reflects relative computation time.

##### Individual model training and testing

For individual models, continuous folds or vocalisations were split into training and test sets, following procedures outlined in the previous section (see Figure 1 for a diagrammatic depiction and Figure 3 for a detailed schematic of each analysis step). First, to obtain the optimal lambda (*λ*) parameter for the mTRF model for each individual participant, models were trained over *λ* values ranging from 10^-7^ to 10^7^ (Jessen et al., 2021), using a ‘leave-one-out’ cross-validation procedure. The cross-validation procedure was implemented using the *mTRFcrossval* function in the mTRF toolbox. Using the data allotted to the training sets only, for each lambda parameter (*λ*), a TRF was calculated for every trial, bar one (left-out) trial, and then averaged over training trials. The predicted speech signal of the left-out trial was then calculated by convolving the EEG response of that trial with the averaged TRF at each channel, which was then compared to the actual speech signal of the left-out trial using Pearson’s correlation (r). The same procedure was iterated over the number of trials available for that participant and the resulting r value for each iteration averaged. This yielded a predictive power estimate (r) of the model for each lambda parameter (*λ*) tested.

The optimal ridge parameter for that participant (i.e. the parameter yielding the model associated with greatest predictive power) was identified (Crosse et al., 2016) and used to train the model on the training set using *mTRFtrain.* This function computes an mTRF model for each training set and then averages across models. Finally, the model was tested using the data left aside in the test set, using *mTRFpredict*: the averaged training model was convolved with the neural response to reconstruct the speech stimulus, and the predictive accuracy between the actual speech signal and that predicted by the model computed via Pearson’s correlation (r). The resulting r value was taken as the overall measure of speech-brain tracking for that participant. Model values were subsequently averaged across participants.

Finally, to examine the extent to which the mTRF models were overfit to the training data, predictive accuracy values during model training and testing were compared. To do so, the predictive accuracy values for the lambda parameter yielding the best predictive accuracy for each participant during cross-validation were compared with the predictive accuracy scores obtained during model testing (Crosse et al., 2021).

##### Generic model training and testing

For the generic models, a model was trained and tested for each participant, by averaging over participant-specific mTRF models across all other participants (see Figure 1 for a diagrammatic depiction and Figure 3 for a detailed schematic of each analysis step). Whilst there are different approaches to training and testing procedures in computing generic models (Crosse et al., 2021; Di Liberto & Lalor, 2017), we follow the procedure outlined by Jessen et al. (2021), which has previously been used to conduct mTRF analyses on data sets recorded from infants.

First, to identify the best *λ* parameter for each participant (using the same parameters as the individual model), an inter-participant cross-validation procedure was conducted. For each *λ* value, for each participant, an mTRF model was estimated using the *mTRFtrain* function in the mTRF toolbox (see above for description). Then, for each participant, a generic model was computed by averaging over the models of all other participants. The predictive accuracy of the averaged mTRF model for the left-out participants’ speech data was computed using the *mTRFpredict* function. This procedure was repeated for every participant at each *λ* value, and the *λ* term yielding the best predictive accuracy for each participant identified. Second, an mTRF model for each participant was computed using the *mTRFtrain* function, inputting the most optimal *λ* value identified for that participant during the cross-validation procedure. To train a generic model for each participant, the models resulting from the optimal *λ* terms were averaged across all other participants and the predictive accuracy of the averaged model tested on the target participants’ speech data, using *mTRFpredict*. Resulting r values were subsequently averaged across participants.

#### 2.2.4. Step 4 - Permutation testing

##### Continuous analysis

To create a random permutation distribution for the continuous models, each participant’s synchronised and downsampled speech stream was randomly paired with another participant’s EEG stream. Similar to the procedures outlined above, the longer of the two data streams was shortened so that both data streams were of the same length.

Corresponding parts of the EEG and speech signals that were set as missing data points in one stream were identified in the other alternate stream and set to missing data points. Each stream was then z scored, and corresponding missing data points set to 0. Exactly the same procedure for computing the continuous folds, and allotting folds to training and test sets, was then repeated in the same way as the main analysis.

A generic or individual mTRF model was then generated and the overall r value computed. This same procedure was repeated 100 times, and r values averaged over the 100 iterations. To test whether the observed values in the actual data differed from chance, a t-test was computed between the observed values and the averaged permutation values. Before conducting the t-test, outliers 1.5 interquartile ranges above the upper quartile and 1.5 interquartile ranges below the lower quartile were excluded (Attaheri et al., 2022).

##### Speech chunking analysis

To create a random permutation distribution, again, each participant’s synchronised and downsampled speech stream was randomly paired with another participant’s EEG stream. Here, each vocalisation extracted from the participant’s synchronised speech stream was paired with a segment of the randomly paired participant’s EEG signal, that matched the length of the vocalisation. Again, where either the speech segment or EEG segment contained any missing data points, this vocalisation was excluded from analysis. To optimise the number of vocalisations included in the permutation distribution, where a segment of the randomly paired EEG data contained missing data points, the algorithm searched a further 3 times for a segment of data of equivalent length but without any missing data values: this number was capped to deal with the fact that some caregivers had some vocalisations that lasted over 1 minute (and therefore had a much higher chance of being paired with EEG data with missing data points). The speech and EEG segments were then z scored and allotted to training and test sets, following exactly same procedures outlined above.

Again, a generic or individual mTRF model was then generated and the overall r value computed. This same procedure was repeated 100 times, and r values averaged over the 100 iterations. To test whether the observed values in the actual data differed from chance a t-test was computed between the observed values and the averaged permutation values, after outlier removal (Attaheri et al., 2022).

## 3. Results

The results section is organised in 3 parts. First, we present descriptive statistics relating to the temporal characteristics of the caregivers’ speech. Second, we present the results of the individual mTRFs, and, finally, the results of the generic mTRF models.

### 3.1. Descriptives

First, we examined the temporal characteristics of caregiver vocal behaviours during naturalistic interaction with their infants. The results of these analyses are presented in Figure 4. Figure 4a shows the proportion of time that caregivers and infants vocalised on their own, as well as the proportion of time they vocalised at the same time (co-vocalisations). This shows that caregivers spent much more time vocalising in comparison to their infants, who vocalised on their own less than 10% of the interaction time. In comparison, caregivers most often vocalised on their own. These findings correspond to our previously reported results, where the start and stop times of caregiver and infant vocalisations were hand-coded in a sub-sample of the data reported here (Phillips et al., 2023).

**Figure 4.**
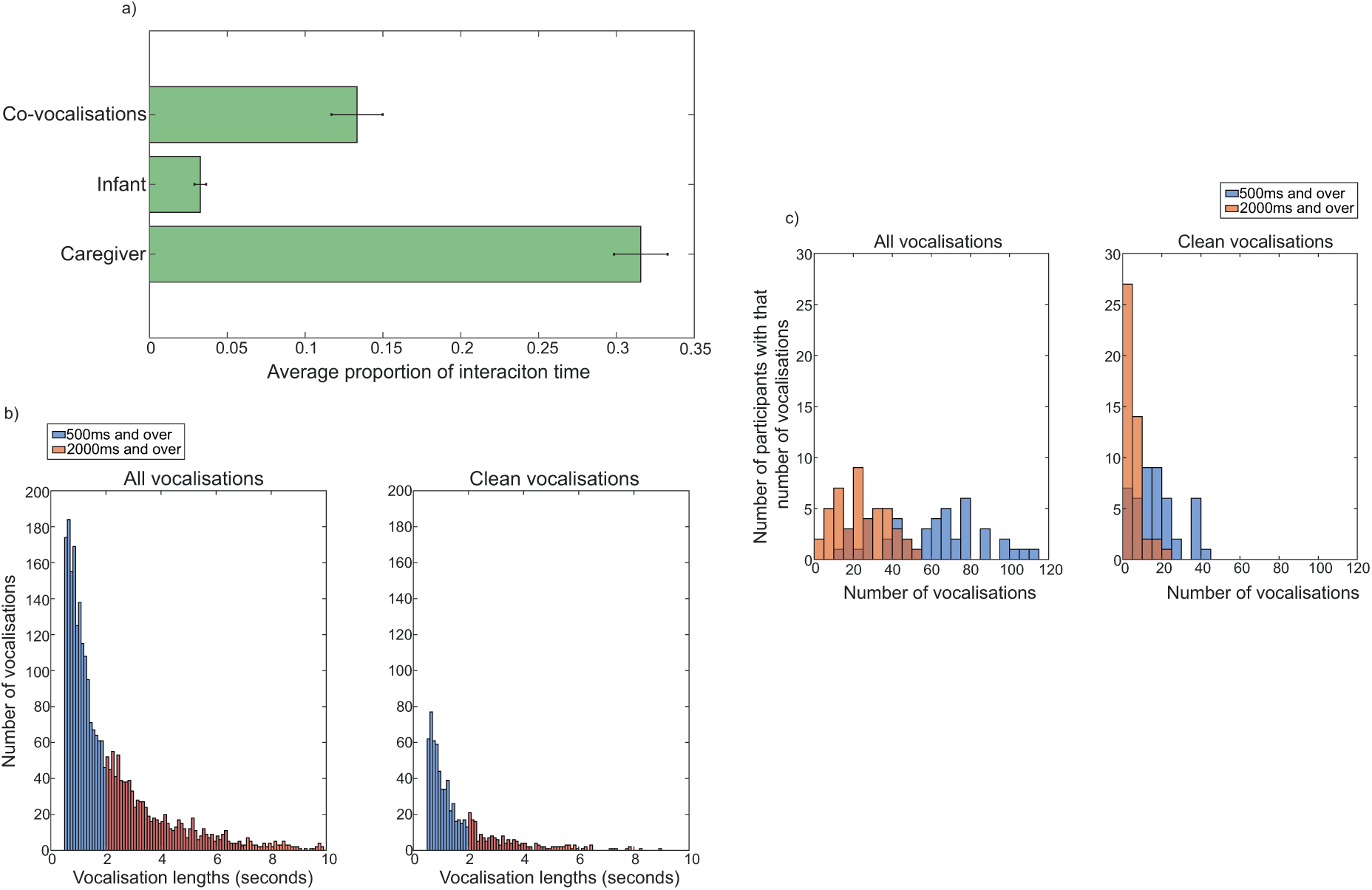
Temporal characteristics of caregiver vocal behaviours. a) Average proportions of time caregivers and infants spent vocalising, and co-vocalising. b) Histogram of the lengths of caregiver vocalisations for all vocalisations (left) and clean vocalisations only (right), over 500ms (blue) and 2000ms (red), after outlier removal. c) Number of caregivers with a given number of vocalisations for all vocalisations (left) and clean vocalisations (right) only lasting over 500ms (blue) and over 2000ms (orange).

Inspection of Figure 4b and 4c shows that the majority of caregiver vocalisations lasted less than 2000ms, with the median vocalisation length for vocalisations lasting over 500ms being 1.51s (IQR=1.24-1.84) and 3.29s (IQR=2.84-3.87) for vocalisations lasting over 2000ms. Corresponding to this, the median number of vocalisations lasting over 2000ms (M=22, IQR=13-34) was much lower compared to those lasting over 500ms (M=62, IQR=40-77). This number was even lower when considering just clean vocalisations over 500ms (M=15, IQR=8-21) and over 2000ms (M=3, IQR=1-7)). Given the particularly low number of vocalisations that were clean and lasted over 2000ms, analyses including these vocalisations only were not conducted. For all vocalisations over 2000ms, 2 participants were removed from analyses because they contributed fewer than 5 vocalisations. For clean vocalisations over 500ms, 7 participants were excluded from analyses for the same reason (see Fig 4c).

### 3.2. Individual mTRF models

In this section, we outline the results of the individual mTRF models obtained for each modulation frequency band, and consider the findings relative to indices of how well the trained model transfers to new data (see Methods (section 2.2.3) for further description).

#### 3.2.1. Continuous individual models

The results of the continuous individual models are presented in Figure 5. The number of outliers removed for each analysis are presented in Table 1. Figure 5a shows the mean predictive accuracy scores across participants for each frequency band. A series of two-tailed paired sample t-tests indicated that only the predictive accuracy values of the delta rate model were significantly higher than the values obtained in the permutation distribution (see Table 1). Testing whether the models differed to chance with Wilcoxon Signed Ranks tests, before outlier removal, yielded a similar pattern of results, with predictive accuracy values significantly above chance for the delta model only (Figure S1).

**Figure 5.**
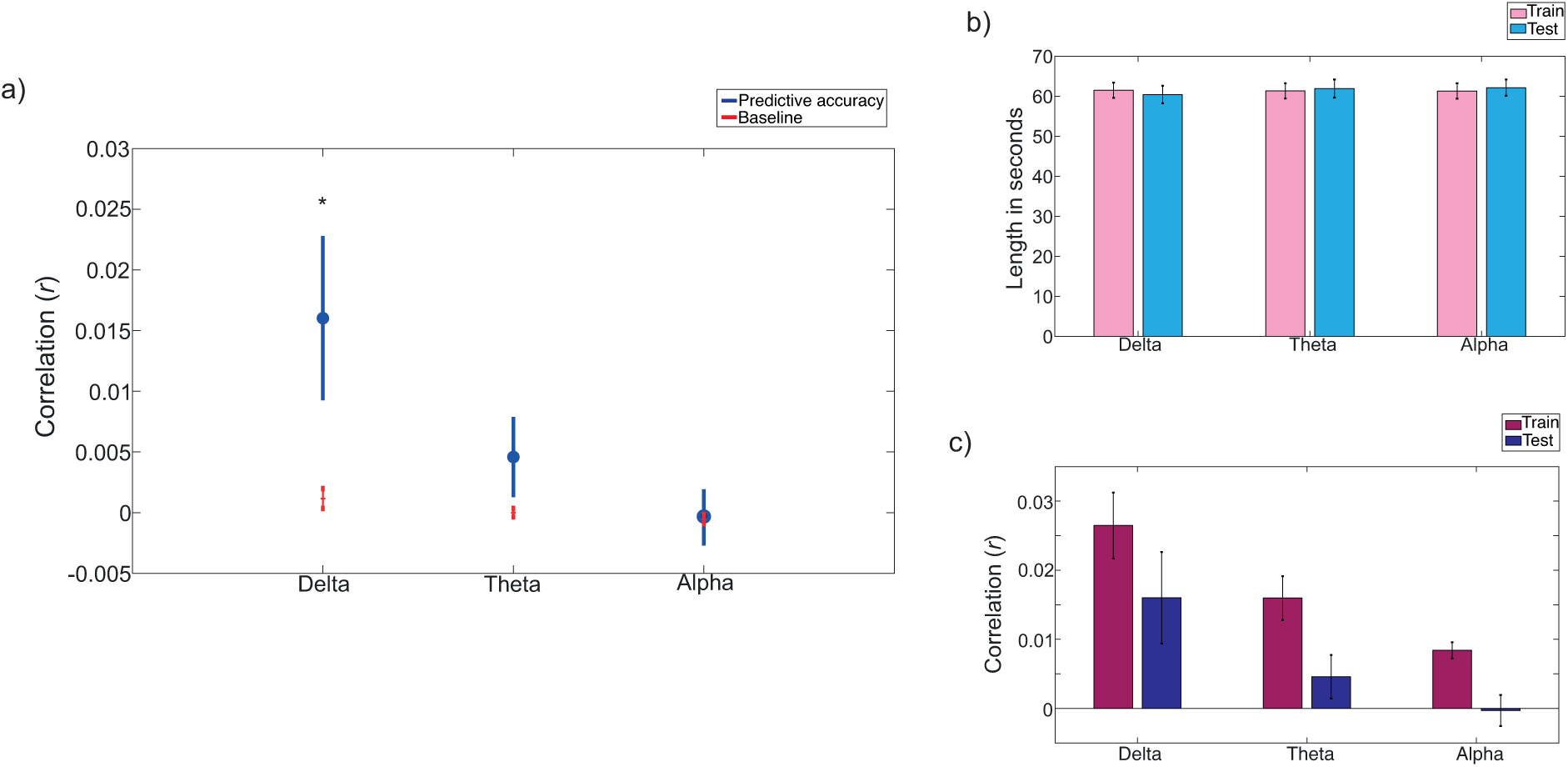
Continuous individual models. a) Grand average predictive accuracy values for each frequency band (Pearson’s r). Blue circles show the mean across participants for each frequency band. Blue lines indicate the SEM of the averaged predictive accuracy values. Red horizontal lines show the averaged permutation values, averaged across permutations and participants. Red vertical lines indicate the SEM. Two-tailed paired sample tests compared the observed r values to chance (*p<0.05). b) Lengths of the folds used in training and testing for each frequency band. Pink bars show the mean lengths of the training sets, and blue bars show the mean lengths of the test sets. Black lines show the SEM. c) Averaged predictive accuracy for training and testing sets. Pink bars show the training sets and blue bars show the test sets. Black lines indicate the SEM.

**Table 1.**
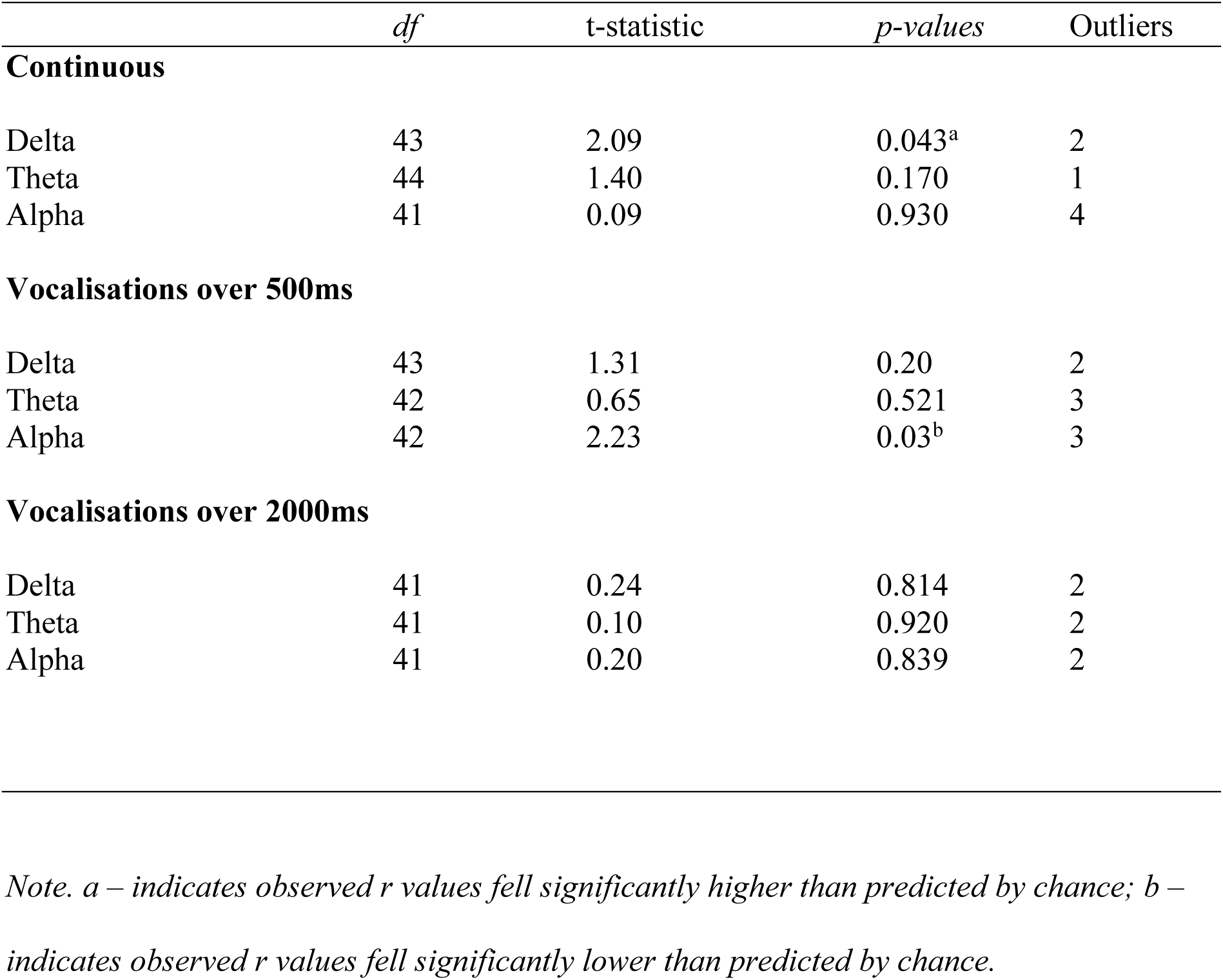
Individual model results of the paired sample t-tests of the difference between observed predictive accuracy scores and the permutation distribution for each frequency band, for each analysis.

Across the different frequency bands, the lengths of the folds included in the training and test sets are similar, and these lengths are consistent across frequency bands (Figure 5b). This is important to consider as the lengths of the folds included in the training and test sets could differ in the continuous models, given that the folds computed for play section 1 could be different in length, compared to those from play section 2, if one interaction was longer than the other (refer to Methods for more detail). Inspection of the error bars (Figure 5b) also reveals little variability between participants, suggesting that the models for each participant were trained and tested on continuous folds of a similar length. The predictive accuracy values computed in model testing, were, however, lower compared to those obtained during cross-validation, for all frequency bands (Figure 5c), which could be an indication that the models are over-fit during model training and therefore do not transfer well to the model at test. That said, for all frequencies, the difference in predictive accuracy values is small (see the Discussion (section 4) for more detailed discussion on this).

#### 3.2.2. Vocalisation individual models

Next, we examined the results of the individual mTRF models computed with individual vocalisations serving as the model folds. First, we tested the performance of individual models, trained and tested on vocalisations lasting 500ms or longer. The results of this analysis are shown in Figure 6. Inspection of Figure 6a shows that, across frequencies, the lengths of the training and test sets were similar, and there was little variability in this across participants. Comparing the predictive accuracy of the models during training and testing (Figure 6b) reveals, that, again, predictive accuracies were much lower for the testing sets compared to the training sets. Relative to Figure 5c, where the data for the continuous individual models is presented, this difference is much greater, and the predictive accuracy (r) values for both training and testing sets are much higher in comparison to those observed where continuous data chunks are used. The results of the individual mTRF model are presented in Figure 6c. A series of paired sample t-tests revealed that mean predictive accuracy values did not fall significantly above their permutation distribution at any frequency band examined (see Table 1 for the results of the t-tests and information on outlier removal).

**Figure 6.**
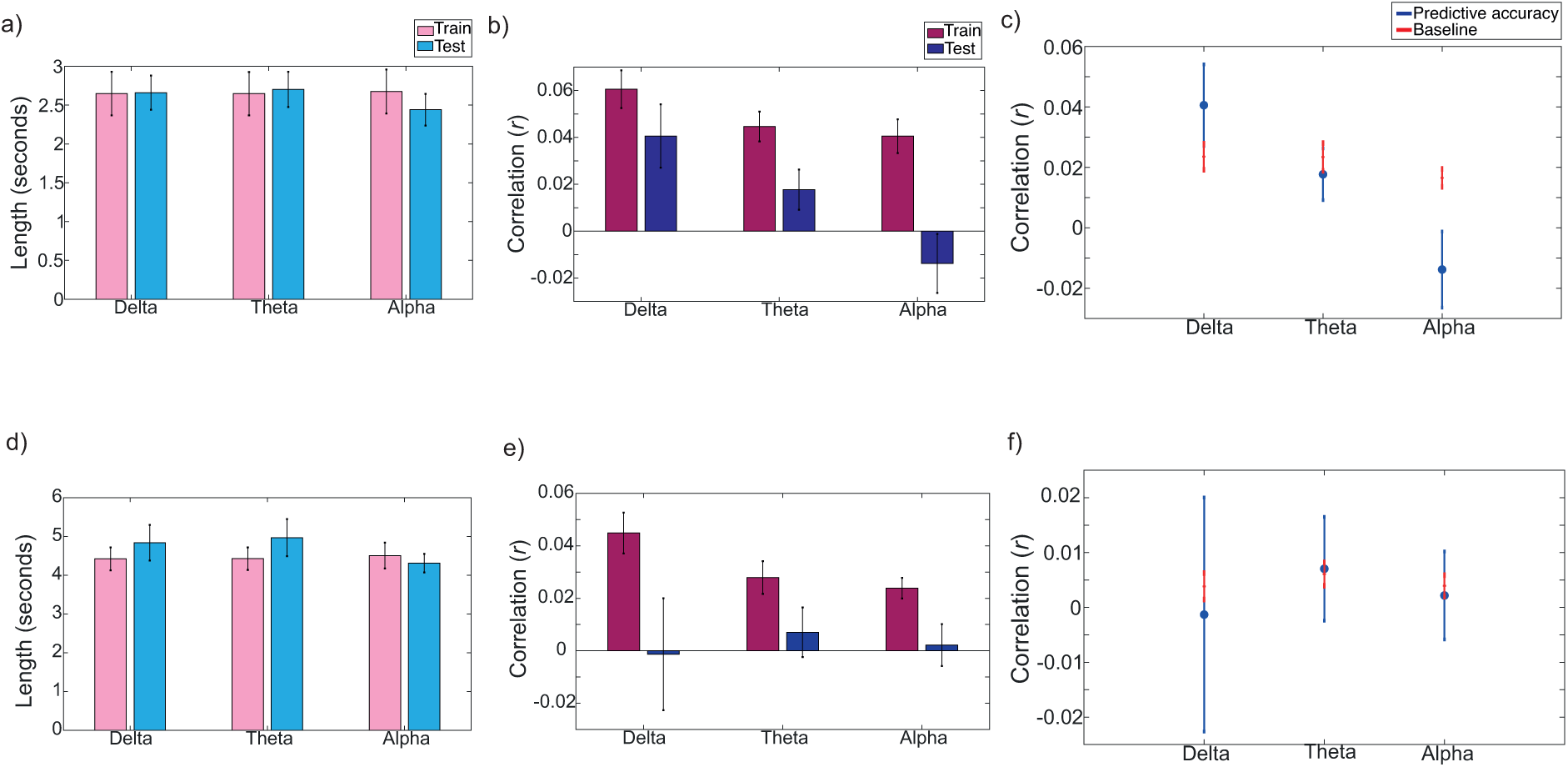
Vocal chunk individual models. First column shows the lengths of the vocalisations included in the training and tests sets for a) vocalisations over 500ms and d) vocalisations over 2000ms. Pink bars show the mean for the training sets and blue bars show the mean for the tests sets. Black lines indicate the SEM. Second column shows the averaged predictive accuracy for training and testing sets for b) vocalisations over 500ms and e) vocalisations over 2000ms. Pink bars show the training sets and blue bars show the testing sets. Black lines indicate the SEM. Last column shows the grand average predictive accuracy values for each frequency band (Pearson’s r), for c) vocalisations over 500ms and f) vocalisations over 2000ms. Blue circles show the mean across participants for each frequency band. Blue lines indicate the SEM of the averaged predictive accuracy values. Red horizontal lines show the averaged permutation values, averaged across permutations and participants. Red vertical lines indicate the SEM. Two-tailed paired sample tests compared the observed r values to chance (*p<0.05).

Next, we examined the performance of the individual models conducted with vocalisations lasting 2000ms or longer (Fig 6d-6f). As mentioned in the Descriptives section (Figure 4b), excluding vocalisations between 500-2000ms led to the inclusion of far fewer vocalisations in the mTRF model. Correspondingly, the lengths of the vocalisations included in the training and test sets are much longer, compared to the analysis using all vocalisations over 500ms (Figure 6d). Comparing the predictive accuracy values in the training and test sets indicated that predictive accuracies at test were substantially lower than those computed during model training (Figure 6e). Though the predictive accuracies, across the models at different frequencies, showed a similar pattern to those obtained in the continuous analysis, correlation values were much lower, and no values were significantly above chance (see Table 1 and Figure 6f). A more detailed discussion of the possible reasons for the pattern of findings reported for these models is included in the Discussion section. Finally, we examined models for clean vocalisations lasting 500ms or longer. The small number of vocalisations entered into these models resulted in seemingly spurious predictive accuracy values. The results of these models are, therefore, reported in the SM (Figure S2 and Table S1).

### 3.3. Generic mTRF models

In this section, we examine the predictive accuracy of the mTRF models yielded by the subject-independent generic models.

#### 3.3.1. Continuous generic models

The results of the continuous generic models are presented in Figure 7a. Similar to the individual models derived from the continuous data, inspection of Figure 7a indicates higher predictive accuracy values at delta and theta frequencies, compared to alpha frequencies.

**Figure 7.**
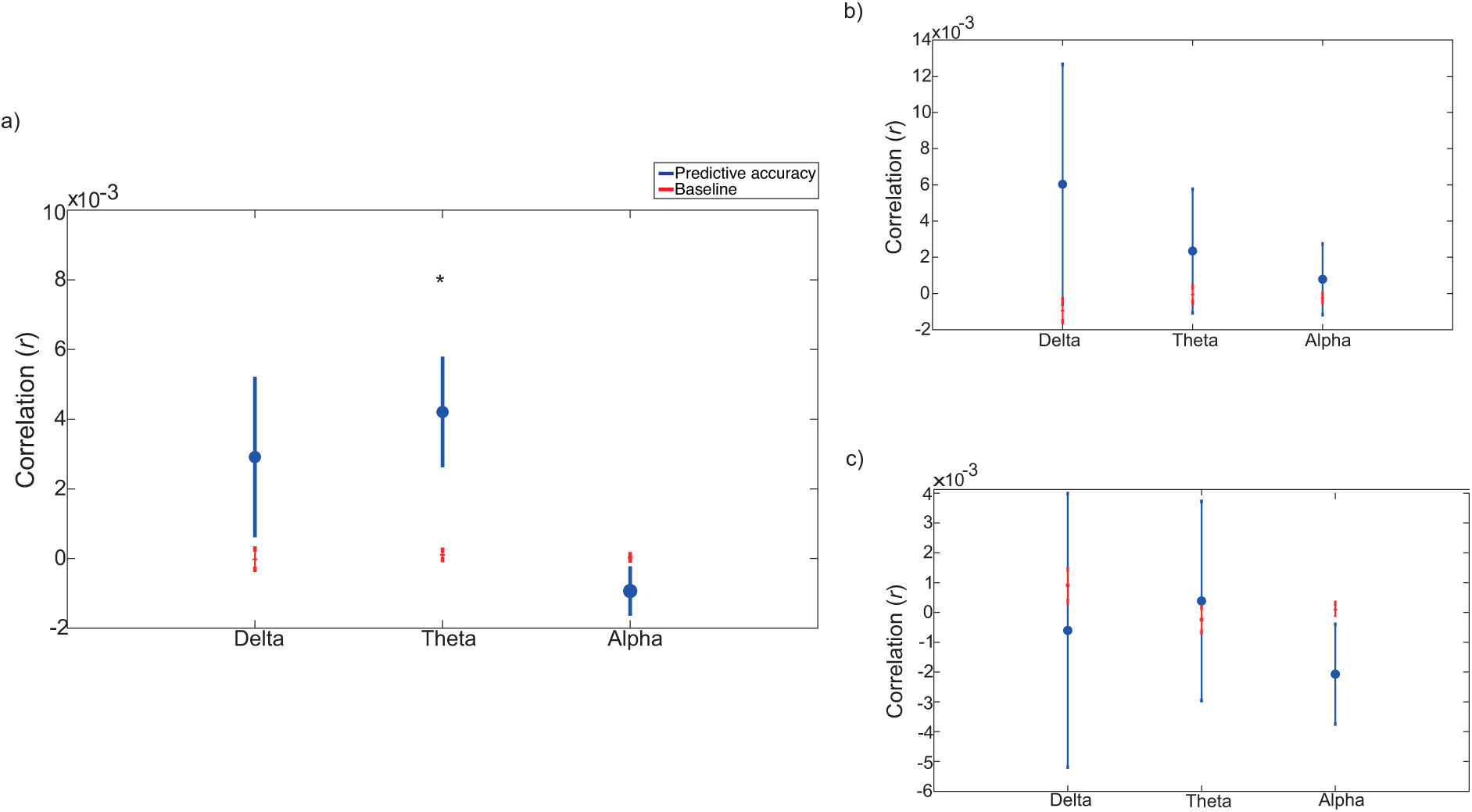
Continuous and vocal chunk generic models. Grand average predictive accuracy values for each frequency band (Pearson’s r) for a) continuous data, b) vocalisations over 500ms and c) vocalisations over 2000ms. Blue circles show the mean across participants for each frequency band. Blue lines indicate the SEM of the averaged predictive accuracy values. Red horizontal lines show the averaged permutation values, averaged across permutations and participants. Red vertical lines indicate the SEM. Two-tailed paired sample tests compared the observed r values to chance (*p<0.05).

Corresponding to this, a series of paired sample t-tests revealed that only the r *-*value of the theta frequency model fell significantly above chance (see Table 2 for the results of the t-tests and information on outlier removal). Of note, however, and in line with previous research with infants using a generic modelling approach (Jessen et al., 2019), the predictive accuracies of the models at all frequency bands are low, and much lower than the individual models (Figure 7a). Comparing the observed values to the permutation distribution using Wilcoxon signed ranks tests, before outlier removal, revealed significant speech-brain tracking at delta and theta frequencies (Figure S3).

**Table 2.** Generic model results of the paired sample t-tests of the difference between observed predictive accuracy scores and the permutation distribution for each frequency band, for each analysis.

|  | *df* | t-statistic | *p-values* | Outliers |
| --- | --- | --- | --- | --- |
| **Continuous** |  |  |  |  |
| Delta | 43 | 1.32 | 0.195 | 2 |
| Theta | 41 | 2.46 | 0.018<sup>a</sup> | 4 |
| Alpha | 40 | 1.35 | 0.184 | 5 |
| **Vocalisations over 500ms** |  |  |  |  |
| Delta | 45 | 1.04 | 0.305 | 0 |
| Theta | 45 | 0.66 | 0.511 | 0 |
| Alpha | 44 | 0.51 | 0.614 | 1 |
| **Vocalisations over 2000ms** |  |  |  |  |
| Delta | 41 | 0.31 | 0.758 | 2 |
| Theta | 42 | 0.18 | 0.857 | 1 |
| Alpha | 40 | 1.20 | 0.236 | 3 |

Note. *a* – indicates observed $r$ values fell significantly higher than predicted by chance; *b* – indicates observed $r$ values fell significantly lower than predicted by chance.

#### 3.3.2. Vocalisation generic models

The generic models computed for vocalisations of differing lengths are reported in Figure 7. For consistency with the reporting of the Individual models, models computed for clean vocalisations lasting over 500ms are reported in the SM (Figure S4 and Table S2). The generic models conducted with vocalisations lasting over 500ms showed a similar pattern to the continuous models, with higher predictive accuracies at theta and delta frequencies, though their predictive accuracies were much lower and more variable (Figure 7b). None of the models at any of the frequency bands investigated differed significantly from the permutation distribution (Figure 7b). Inspection of Figure 7c shows that where models included vocalisations only lasting over 2000ms, predictive accuracies were even lower, with even greater variability, and this pattern was the same for models conducted with clean vocalisations lasting over 500ms (Figure S4). Again, r values for these models did not differ significantly to the permutation distribution (for further information on statistical testing see Table 2).

## 4. Discussion

In this study we examined the applicability of continuous methods of analysis for assessing speech-brain tracking during naturalistic caregiver-infant table-top play. Whilst these methods are increasingly being used in adult and infant populations to assess speech-brain tracking relative to pre-recorded stimuli, no previous work has applied these methods to continuous, free-flowing interactions. Investigating neural tracking of the speech signal in naturalistic contexts is crucial to our understanding of how infants process the phonological information of speech during everyday communication (Attaheri et al., 2022; Leong et al., 2017), as well as the association of these micro-processes to the ongoing behavioural dynamics of early social interactions. We conducted analyses similar to those outlined in Attaheri et al. (2022), employing a backwards modelling mTRF approach, to model the relationship between the infants’ continuous neural response and their caregiver’s speech signal. To maximise the amount of data included in the models, we first conducted training and testing procedures on continuous data segments of the interaction, and then on individual vocalisation chunks, after excluding vocalisations below certain minimum lengths. To train the models, both individual and generic methods were employed. For each model, we examine differences between predictive accuracy values computed during model training and model testing, to check the extent of over-fitting to the training data.

For the individual models, where continuous folds were entered into the model (i.e. where interactions for each participants were segmented into 10 chunks and submitted to the mTRF model; see section 2.2.3), significant speech-brain tracking was observed at delta frequencies, but not at theta or alpha frequencies (Figure 5a), corresponding to the findings of Attaheri et al. (2022). For these models, predictive accuracy values were greater during model training compared to testing, but this difference was small. This suggests that, where individual mTRF models are trained and tested on continuous periods of data obtained by dividing the interactions up into folds of equal lengths, model over-fitting to the training data is low (Crosse et al., 2021).

In the models that were trained and tested on individual vocalisation chunks of 500ms or more, the predictive accuracy values showed the same overall pattern but none of the values fell significantly above chance. In comparison to the models trained on continuous data, the difference in the predictive accuracy values acquired in model training and testing was greater (Figure 6b,c), suggesting that model overfitting to the training data was more problematic in this analysis compared to the continuous training method. Greater over-fitting to the training set likely reflects the much smaller amount of data entered into the model, compared to those trained on continuous data folds, that utilised all audio data occurring across the entire interaction.

Also of note, the predictive accuracy values for the delta and theta models are in fact higher in comparison to the models trained on the continuous data sets; but, unlike the continuous models, the permutation distributions fall above 0, resulting in a non-significant difference between observed and averaged permutation models. The inflated pattern of predictive accuracy values, across observed and permuted data sets is likely a result of the short data folds entered into the models trained on individual vocalisations lasting over 500ms, which predominantly included data chunks between 500 and 2000ms in length (see Figure 4). This supports the conclusions of Crosse et al. (2021) who highlighted that using window sizes of less than 10s when computing mTRF models in sliding windows can result in spuriously high and low predictive accuracy values, as well as decreases in speech-tracking estimates.

Corroborating this interpretation, where far fewer and much shorter vocalisations were included in model training and testing, in models trained only on *clean vocalisations lasting over 500ms* (i.e. there were fewer clean vocalisations and these also tended to be shorter, as longer vocalisations were more likely to pick up noise), model estimates fell well below 0, yielding findings that are likely to be artifactual across all frequencies (see Figure S4). Where longer but fewer vocalisations were entered into model training and testing (i.e. models including *only clean vocalisations over 2000ms*), predictive accuracy values were close to 0 (Figure 6f), and a comparison of training and test estimates (Figure 6e) indicates model overfitting to the training data; most likely this reflects the decreased amount of data entered into this model, in comparison to the continuous (Figure 5b) and 500ms models (Figure 6a).

Overall, the results of the individual models suggest that, with a subject-dependent training procedure, most reliable speech tracking values are obtained where the interaction is divided up into continuous folds (i.e. where the entire interaction is split into equal-length folds, irrespective of when the caregiver is speaking), and, therefore, where the greatest amount of data is used in model training.

Given the issues of data quality and quantity inherent to naturalistic data recordings, in the second part of this paper we computed mTRF models using generic training procedures which train models by utilising all the available data for each participant (Jessen et al., 2021). Whilst generic models reduce the likelihood of overfitting with smaller and more noisy data sets, the predictive power of these models is often lower in being trained across participants, with data sets of variable quality (Jessen et al., 2021). The results of the generic models showed that, in both types of training procedures (continuous inputs and vocalisation chunks of variable lengths) the predictive accuracy values were very close to 0 and much more variable in comparison to previous studies of speech-brain tracking conducted with infants (Jessen, 2019; Jessen et al., 2021). Significant speech-brain tracking was observed at theta frequencies where the models were trained on continuous data folds, and the delta model also fell significantly above the permutation distribution before outlier removal (Figure S3). The particularly low predictive accuracy scores obtained for the generic models, however, precludes any reliable interpretations of speech-brain tracking from these models, for any frequency band examined (Crosse et al., 2021).

The low and variable predictive accuracy values obtained with the generic models are likely driven by the fact that, in our naturalistic interactions, infants were interacting with their caregiver and therefore listening to different speech signals. This stands in contrast to controlled paradigms where each infant listens to the same stimulus (e.g. Jessen, 2019). This will drive variation in model fitting in two ways: the amount of noise in the speech signal is likely to vary substantially across participants, and aspects of the noise and characteristics of the speech signal could also drive differences in how infants track the amplitude envelope. Furthermore, whereas in controlled paradigms infants are often seated on their caregiver’s lap and encouraged to stay still as much as possible throughout the recording, in naturalistic interactions, infants are free-moving, meaning that the amount of movement and, therefore, the amount of movement-related artefact affecting the EEG signal will also vary substantially across participants (Noreika et al., 2020). The extent to which variability in the noise of either the EEG or speech signal vs. the variability in the spectral and temporal qualities of the caregivers’ speech signal, and individual differences in how infants track their caregiver’s speech signal, should be a key focus for future research. A more detailed discussion on how this could be conducted is given below.

Overall, the results of the individual and generic models indicate that the use of individual training procedures, conducted on continuous segments of the interaction, may be most meaningful in examining speech-brain tracking during naturalistic social interactions. The difference between the predictive accuracy scores for training and test sets was smallest for the continuous data models, compared to the vocal chunk models, and variability in predictive accuracy values was also low. Computing models using generic training procedures associated with particularly low speech-tracking values.

Given that the inputs to the mTRF model with continuous analysis procedures contained frequent segments of noise unrelated to the speech of the caregiver, including toy clacks and infant vocalisations, it will be important to conduct further analyses to support our assumption that the neural tracking findings observed here are driven by alignment between the infants’ EEG activity and modulatory changes in the adults’ speech. The mTRF modelling approach is particularly robust to the inclusion of missing data points (even up to 30-40%; Jessen et al., 2021). One option would therefore be to use a continuous chunking approach and insert 0 values at points in the amplitude envelope that the caregivers are not speaking. That said, inspection of Figure 4a reveals that, in our data sets, caregivers vocalised on their own for an average of 35% of the interaction, meaning that, even with the robustness of the mTRF models to missing data points, this analysis would not be possible. To examine this in more detail, therefore, in future work, it will be important to apply mTRF modelling procedures to naturalistic interactions where the caregiver speaks more often and the interaction is less affected background noise (e.g. caregiver puppet shows with their infant). Another particularly important avenue for future work examining speech-brain tracking in naturalistic contexts is in testing the effects of different aspects of noise in both the EEG and speech signals on the performance of the mTRF analysis, with infant populations.

Nevertheless, the results of the continuous individual models are consistent with the prediction that delta rate modulations play a particularly important role in facilitating infants’ early ability to parse the speech signal (Leong et al., 2017; Leong & Goswami, 2015). Leong et al., (2017), for example, showed that rather than containing most power at theta rate modulations (reported for ADS), IDS is associated with peak modulation frequencies at delta-rate modulations, with the peak frequency increasing from 7-11 months. The results of the individual models also correspond to Attaheri et al.’s (2022) findings, where speech-brain tracking was found to be significantly above chance at both delta and theta frequencies but not alpha frequencies: and tracking at delta rate frequencies was significantly higher in comparison to theta frequencies at all 3 ages examined. One possible explanation for why we found speech-brain tracking at delta frequencies only could be because the Attaheri experiment used nursery rhyme stimuli, where a British female speaker melodically sung or chanted nursery rhymes, in time with a metronome at 120 bpm. This speech would be inherently more rhythmic compared to the spoken IDS of the caregivers in our study, likely resulting in a higher proportion of stressed syllables (Leong, Kalashnikova, et al., 2017; Leong & Goswami, 2015).

It is interesting and potentially important to note, however, that, similar to the findings reported by Attaheri et al. (2022), the permuted predictive accuracy values reported in Figure 5, generated with randomly paired, unsynchronised data streams, show the same pattern as the real data: i.e. higher values are observed at lower EEG frequencies. This could indicate that the pattern of findings seen in the real data, at least in part, reflects aspects of the underlying structure of the EEG data. The variability of the amplitude values at lower frequencies, for example, will be much greater, compared to the higher frequencies (Cohen, 2014). This is likely to result in a better regression fit between the EEG data and the speech signal (which was held constant in models computed across all EEG frequencies; Kamel & Abonazel, 2023). This fact alone could also explain the increased variability of predictive accuracy values of the models computed for lower, compared to higher EEG frequency bands (see Figure 5). Recent work has also shown that infants’ EEG signals at higher frequencies (particularly > 12Hz) are more affected by movement related artefacts, including limb, jaw and head movements, compared to lower frequencies (Georgieva et al., 2020). The pattern of observed and permuted predictive accuracy values across frequency bands could, potentially, therefore, also be driven by increases in signal to noise ratio at lower EEG frequencies, meaning that the associations between infant oscillatory neural activity and the speech signal are more easily detected. That said, despite the overall pattern of findings across frequencies being the same for the observed and permuted data, the delta-rate prediction accuracies do fall significantly above their permutation distribution, whereas the theta and alpha values do not.

It will be important for future work to develop a more mechanistic framework for understanding how infants attune to the fine-grained temporal dynamics of their caregivers’ speech at different modulation frequencies. This will require a combination of structured, semi-structured and naturalistic paradigms, where infant neural activity is recorded in response to continuous speech streams. To provide further support for the idea that delta-rate tracking is particularly important in early infancy, experiments examining infant neural tracking of continuous pre-recorded speech, where the sharpness of temporal fluctuations in the amplitude envelope at lower modulation frequencies are degraded should be conducted. Neural tracking of only non-degraded, compared to degraded speech, which has previously been shown in adults (Doelling et al., 2014), would provide more clear support for the particular importance of speech-brain tracking at lower modulation frequencies in early infancy.

Given that delta rate modulations in speech drive the perception of stressed syllables, which are thought particularly important to early word segmentation, (Leong et al., 2017; Leong & Goswami, 2015), further analysis of infant neural activity relative to stressed syllables in naturalistic nursery rhyme and free-flowing infant-directed speech should also be a key focus. Rather than the continuous methods of analysis reported here, such investigations could employ event-locked analysis methods to examine phase synchronisation between the amplitude envelope of the caregivers’ speech signal and infant EEG activity at the onsets of stressed syllables in caregiver speech (Ní Choisdealbha et al., 2023; Power et al., 2013). In naturalistic free-play contexts, like the study reported here, the role of toy clacks, infant vocalisations, and other non-vocal interactive noises in disrupting the stress-timed patterning of caregiver speech, and the effect this has on infants’ neural response to delta rate temporal modulations, should also be examined. Further to this, future behavioural and neurophysiological paradigms should examine the associations between the temporal modulations of infant-directed speech, and other perceptually salient aspects of the caregivers’ speech signal, such as fluctuations in fundamental frequency, as well as its higher-level semantic properties (Di Liberto et al., 2023; Legendre et al., 2019; Räsänen et al., 2018; Weineck et al., 2022). This work will support a more holistic understanding of the features of naturalistic linguistic inputs that infants begin to use to make online predictions about speech and the co-occurring behaviours of an interacting partner (Tan et al., 2022)

Overall, this paper has demonstrated the applicability of continuous modelling approaches to examining speech-brain tracking by infants to their own caregivers’ speech signal during naturalistic, free-flowing interactions. The results are important in showing that the finding of significant speech-brain tracking in early infancy to controlled, often purposefully rhythmic stimuli, in experimental paradigms transfers to everyday, naturalistic interactions.

Developing these continuous methods further, in combination with a detailed examination of the precise mechanisms that drive speech-brain tracking in infancy, will open up new avenues for investigating how early attunement to the temporal modulations in speech associates with the joint attention processes of real-time interactions (e.g. mutual gaze and shared attention towards objects), as well as the timing of caregiver object labelling, and the higher-level semantic complexities of their inputs (Nencheva & Lew-Williams, 2022).

## Acknowledgements

Thank you to Navsheen Kaur and Glenda Garcia for help with data coding. Thanks to Trinh Nguyen for reading and commenting on earlier versions of this manuscript. Thank you to members of the UEL BabyDev Lab for comments and discussions on earlier drafts of this manuscript, and to all participating children and caregivers.

## Author Contributions

EAMP designed experiment; collected data; performed analyses; wrote manuscript; LG, PL and JI co-wrote analyses; commented on manuscript; IMH collected data; contributed analyses; MW collected data; SVW secured funding; co-wrote analyses.

## Data and Materials Availability

Due to the personally identifiable nature of our data (video and audio recordings of infants), permission to access the data will be given only by contacting the first author, EAMP, direct via email.

## Supplementary material

### Results

#### Figures

**Figure S1.**
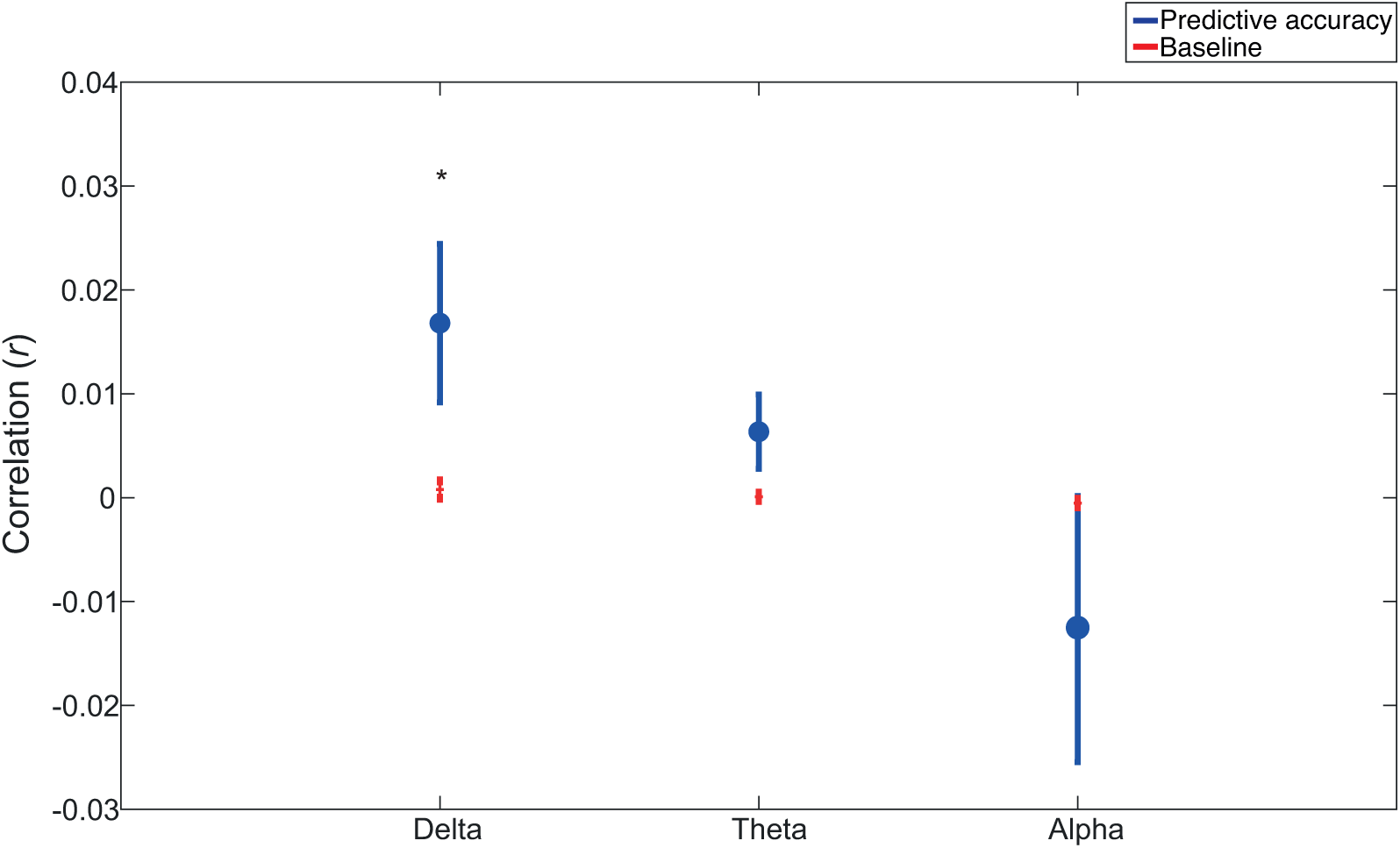
Grand average predictive accuracy values for individual continuous models before outlier removal, for each frequency band (Pearson’s r). Blue circles show the mean across participants for each frequency band. Blue lines indicate the SEM of the averaged predictive accuracy values. Red horizontal lines show the averaged permutation values, averaged across permutations and participants. Red vertical lines indicate the SEM. Wilcoxon Signed Rank tests compared the observed r values to chance (*p<0.05).

**Figure S2.**
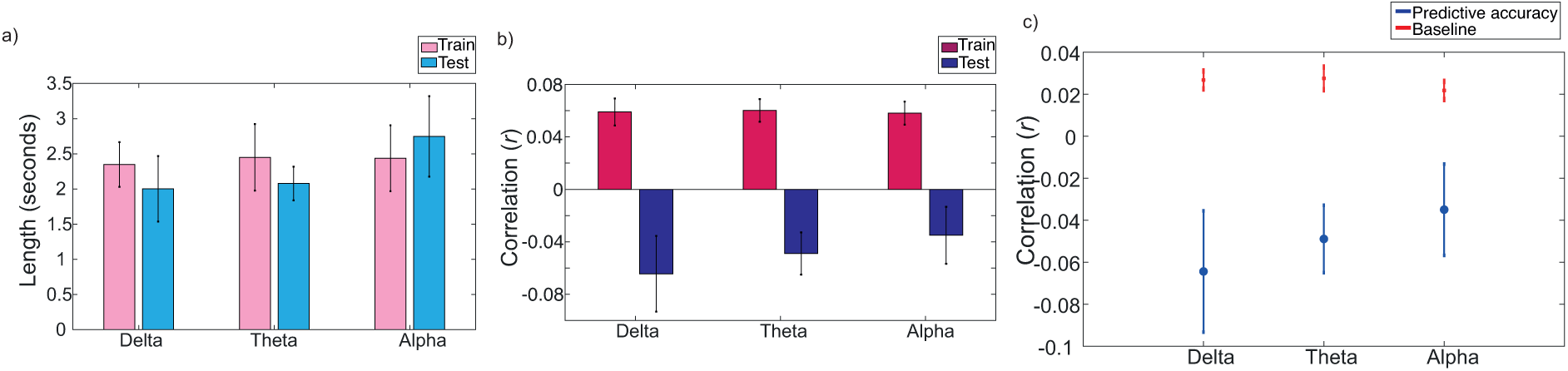
Vocal chunk individual models for clean vocalisations over 500ms in length. a) Lengths of the vocalisations included in the training and tests sets, b) averaged predictive accuracy for training and testing sets c) grand average predictive accuracy values for each frequency band (Pearson’s r). Blue lines indicate the SEM of the averaged predictive accuracy values. Red horizontal lines show the averaged permutation values, averaged across permutations and participants. Red vertical lines indicate the SEM. Two-tailed paired sample tests compared the observed r values to chance (*p<0.05).

**Figure S3.**
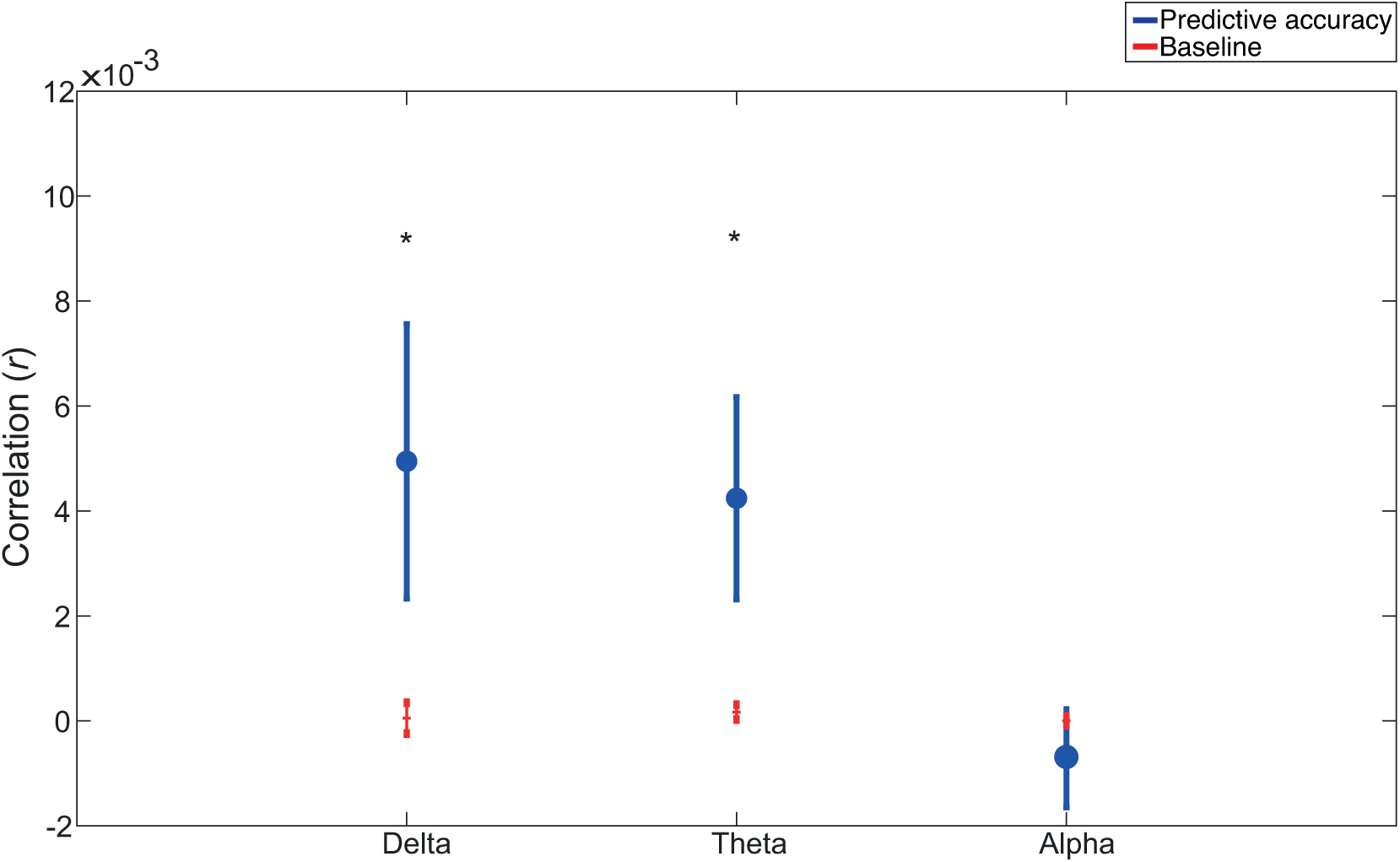
Grand average predictive accuracy values for generic continuous models before outlier removal, for each frequency band (Pearson’s r). Blue circles show the mean across participants for each frequency band. Blue lines indicate the SEM of the averaged predictive accuracy values. Red horizontal lines show the averaged permutation values, averaged across permutations and participants. Red vertical lines indicate the SEM. Wilcoxon Signed Rank tests compared the observed r values to chance (*p<0.05).

**Figure S4.**
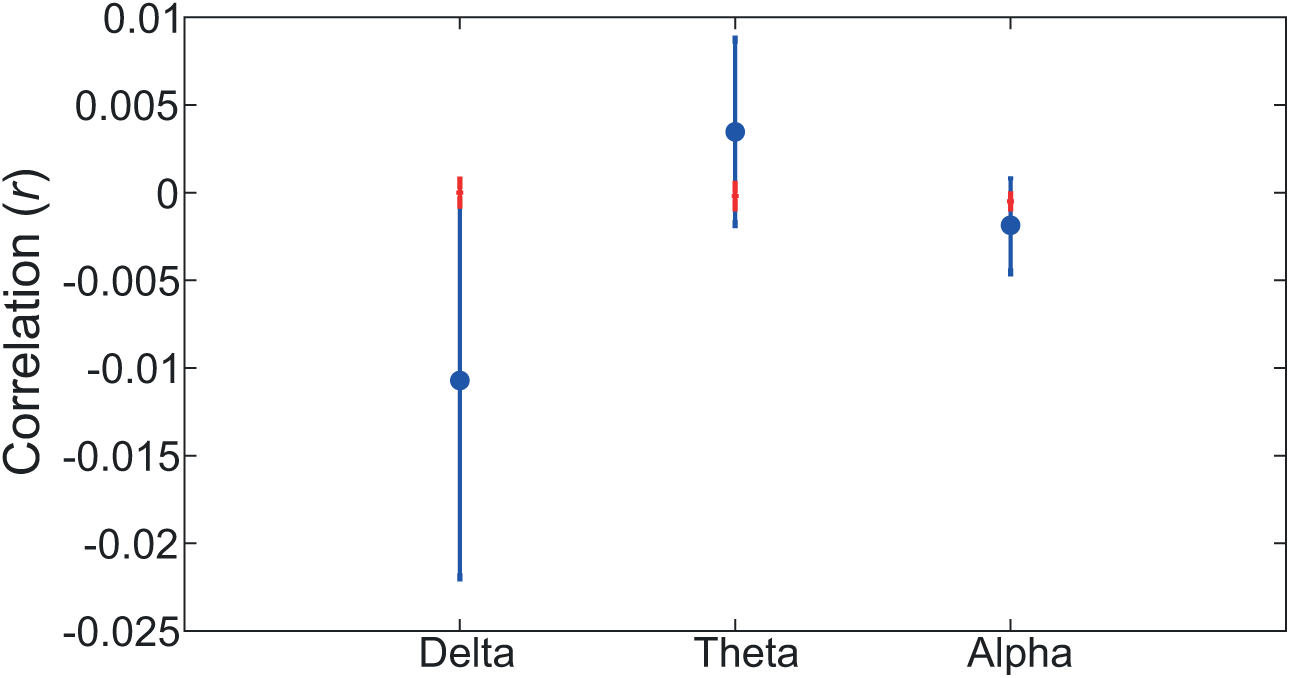
Vocal chunk generic models. Grand average predictive accuracy values for clean vocalisations over 500ms. Blue circles show the mean across participants for each frequency band.. Blue lines indicate the SEM of the averaged predictive accuracy values. Red horizontal lines show the averaged permutation values, averaged across permutations and participants. Red vertical lines indicate the SEM. Two-tailed paired sample tests compared the observed r values to chance (*p<0.05).

#### Tables

**Table S1.** Individual model results of the paired sample t-tests of the difference between observed predictive accuracy scores and the permutation distribution for each frequency band, for clean vocalisations lasting over 500ms only.

|  | *df* | t-statistic | *p-values* | Outliers |
| --- | --- | --- | --- | --- |
| **Clean vocalisations over 500ms** |  |  |  |  |
| Delta | 37 | 3.07 | 0.004 | 1 |
| Theta | 36 | 4.41 | <0.001 | 2 |
| Alpha | 33 | 2.36 | 0.0240 | 5 |

**Table S2.** Generic model results of the paired sample t-tests of the difference between observed predictive accuracy scores and the permutation distribution for each frequency band, for clean vocalisations lasting over 500ms only.

|  | *df* | t-statistic | *p-values* | Outliers |
| --- | --- | --- | --- | --- |
| **Clean Vocalisations over 500ms** |  |  |  |  |
| Delta | 37 | 0.93 | 0.360 | 1 |
| Theta | 38 | 0.70 | 0.485 | 0 |
| Alpha | 34 | 0.47 | 0.638 | 4 |

